# Whole brain delivery of an instability-prone *Mecp2* transgene improves behavioral and molecular pathological defects in mouse models of Rett syndrome

**DOI:** 10.1101/798793

**Authors:** Mirko Luoni, Serena Giannelli, Marzia Indrigo, Antonio Niro, Luca Massimino, Angelo Iannielli, Laura Passeri, Fabio Russo, Giuseppe Morabito, Piera Calamita, Silvia Gregori, Benjamin Deverman, Vania Broccoli

**Affiliations:** Stem Cell and Neurogenesis Unit, Division of Neuroscience, San Raffaele Scientific Institute, 20132 Milan, Italy; CNR Institute of Neuroscience, 20129 Milan, Italy; San Raffaele Telethon Institute for Gene Therapy (SR-Tiget), San Raffaele Scientific Institute IRCCS, Via Olgettina, 58, 20132, Milan, Italy; National Institute of Molecular Genetics (INGM), Milan 20122, Italy; Stanley Center for Psychiatric Research at Broad Institute, Cambridge, MA, USA

## Abstract

Rett syndrome (RTT) is an incurable neurodevelopmental disorder caused by mutations in the gene encoding for methyl-CpG binding-protein 2 (MeCP2). Gene therapy for this disease presents inherent hurdles since *MECP2* is expressed throughout the brain and its duplication leads to severe neurological conditions as well. However, the recent introduction of AAV-PHP.eB, an engineered capsid with an unprecedented efficiency in crossing the blood-brain barrier upon intravenous injection, has provided an invaluable vehicle for gene transfer in the mouse nervous system. Herein, we use AAV-PHP.eB to deliver an instability-prone *Mecp2* (*iMecp2*) transgene cassette which, increasing RNA destabilization and inefficient protein translation of the viral *Mecp2* transgene, limits supraphysiological Mecp2 protein levels in transduced neural tissues. Intravenous injections of the PHP.eB-*iMecp2* virus in symptomatic male and female *Mecp2* mutant mice significantly ameliorated the disease progression with improved locomotor activity, coordination, lifespan and normalization of altered gene expression and mTOR signaling. Remarkably, PHP.eB-*iMecp2* administration did not result in severe toxicity effects either in female *Mecp2* mutant or in wild-type animals. In contrast, we observed a strong immune response to the transgene in treated male *Mecp2* mutant mice that was overcome by immunosuppression. Overall, PHP.eB-mediated delivery of the *iMecp2* cassette provided widespread and efficient gene transfer maintaining physiological Mecp2 protein levels in the brain. This combination defines a novel viral system with significant therapeutic efficacy and increased safety which can contribute to overcome the hurdles that are delaying clinical applications of gene therapy for RTT.

**One Sentence Summary:** Global brain transduction of the instability-prone *Mecp2* transgene by systemic AAV-PHP.eB administration is both safe and effective in protecting male and female *Mecp2* mutant mice from the RTT disease phenotype.

## Introduction

Rett syndrome (RTT) is a severe neurological disorder and second cause of intellectual disabilities in girls. RTT is distinguished by a period of 6-12 month of overtly normal development followed, then, by a rapid regression with the loss of the purposeful motor skills and the onset of repetitive and autistic behaviors^1,2^. In the vast majority of cases RTT is caused by loss-of-function mutations in the *MECP2* gene, which encodes for the methyl-CpG binding protein 2 (MeCP2), a global chromatin regulator highly expressed in neurons^3^. Neuropathological studies have shown that RTT brains exhibit abnormal neuronal morphology, but no sign of neuronal death. *Mecp2*-deficient mice recapitulate key neurological deficits observed in RTT patients offering an invaluable model where to investigate pathological mechanisms as well as test innovative therapies^4^. Similar to human patients, RTT mice show microcephaly without gross neuropathological changes or neurodegeneration. In contrast, recent studies have convincingly revealed that the *Mecp2* loss significantly alters neuronal activity leading to a progressive imbalance of the excitatory-inhibitory synaptic activity across the brain with divergent modalities occurring between different circuits and regions of the brain^5,6^. In 2007, a seminal study by the Adrian Bird’s group demonstrated that the RTT pathological phenotype can be significantly reversed in mice by re-activating the *Mecp2* even at advanced disease stages^7^. In fact, genetic reactivation of *Mecp2* in more than 70% of the neurons in adult mice normalized brain morphology and significantly improved several sensory-motor dysfunctions^8^. These findings provide a strong evidence that Mecp2 is a key factor in maintaining full neurological function during adulthood. Consistently, multiple pathological manifestations exhibited by adult mutant RTT mice can be fully recapitulated by deleting *Mecp2* exclusively in adulthood^9–11^.

Despite the proven genetic reversibility of the RTT disease phenotype in mice, translational treatments aiming at curing the disease or some of its neurological symptoms have not been successful yet. In fact, MeCP2 is a global determinant of the neural chromatin structure and is a pervasive regulator of gene expression in brain cells and, thus, it remains challenging to target a single MeCP2 downstream pathway to obtain a substantial therapeutic benefit^1,2^. The inherent monogenic nature of RTT makes gene therapy a strong translational option for this disease. However, *MECP2* gene duplication in humans is responsible for a serious and clinically distinguished neurodevelopmental disorder. Affected males present with early hypotonia, limb spasticity and severe intellectual disability^12,13^. Thus, a successful gene therapy for RTT has to deliver the correct MeCP2 dosage in a tight range that overlaps with endogenous levels. Different studies have recently provided encouraging results showing that intravenous administration of an Adeno-associated virus serotype 9 (AAV9) expressing the wild-type (WT) *MECP2* attenuated neurological dysfunctions and extended lifespan in RTT mice^14–17^. However, the AAV9 does not cross efficiently the blood-brain barrier in adult mice while having a preferential tropism for peripheral organs. In fact, intravenous delivery of high doses of AAV9 can lead to severe liver toxicity and sudden death due to ectopic MeCP2 expression^15^. Moreover, the limited brain transduction obtained in these studies was not sufficient to determine a correlation between the viral dose, transduction efficiency and therapeutic benefits. Finally, it remains to be determined whether a gene therapy approach is capable to rescue molecular dysfunctions and transcriptional alterations caused by loss of *Mecp2* in the adult brain.

Recently, novel AAV9 synthetic variants have been generated through selected mutagenesis of the capsid proteins to enhance gene transfer in the brain after viral systemic delivery. In particular, using a cell type-specific capsid selection screening, the novel engineered capsid AAV-PHP.B has been developed through a 7-amino-acid insertion in the AAV9 VP1 capsid protein^18^. The AAV-PHP.B viral particles are able to permeate the blood-brain barrier of adult mice and diffuse throughout the neural parenchyma transducing with high efficiency both neurons and glia^18,19^. A subsequent enhanced version has been isolated, named AAV-PHP.eB (PHP.eB in short), with increasing efficiency in brain targeting, thus, requiring lower doses of virus for high neuronal transduction^20^. We sought to use the PHP.eB as viral platform to sustain a global and efficient *Mecp2* brain transduction in adult mutant and WT mice. This system provided us with a flexible and potent tool to determine the efficacy and safety of diffuse *Mecp2* gene transfer in the adult mouse brain. Herein, using PHP.eB transductions in neuronal cultures, we characterized the viral gene expression cassette in all its elements to define its overall transcripts instability and limited translational efficiency. For this reason, the use of a constitutive strong promoter resulted more suitable to restore Mecp2 protein levels in a physiological range and limit viral overload, without compromising its disease rescue efficacy. Indeed, we obtained a robust phenotypic amelioration both in male and female *Mecp2* mutant mice, without severe toxicity effects in wild-type animals. In addition, we identified and characterized a strong immune response to the exogenous Mecp2 in male mutant mice, that severely affected the life-span of treated animals. This limitation was overcome by chronic immunosuppression that granted a life-span extension of up to nine months.

## Results

### Designing an instable *Mecp2* transgene cassette with reduced translation efficiency

Given that the therapeutic range of *Mecp2* gene expression is very limited, earlier studies employed a viral cassette with a minimal *Mecp2* promoter with the attempt to reconstitute endogenous gene expression levels. However, the regulatory regions of *Mecp2* remain poorly characterized and the use of self-complementary AAV9 with a limited cargo capacity has obliged to include only very short fragments of the proximal promoter^15–17,21^. With these conditions, it remains very unlikely to recapitulate the endogenous *Mecp2* expression pattern, whereas the risk of insufficient expression become higher. More generally, the lack of a system to compare exogenous with endogenous levels of MeCP2 is preventing the optimization of the viral transgene cassette. Herein, we overcame these limitations by setting a novel validation assay to compare the transcriptional efficiency of different viral *Mecp2* transgene cassettes directly in neuronal cultures. We initially confirmed that more than 90% of primary mouse neurons can be transduced with the PHP.eB virus. Then, we generated a transgene cassette with the *Mecp2_e1* isoform including the coding sequence and a short proximal 3’UTR (3’pUTR, ∼200bp). This *Mecp2* transcript occurs naturally in embryonic stem cells and various tissues, but during development of the neural system this form is progressively overcome by transcripts with longer 3’-UTR (8,6 Kb)^22^. *Mecp2* transgene sequence carrying an N-terminal V5 tag was driven by two types of promoter (Fig. 1a). We opted to use the chicken-β-actin promoter, which has been extensively used in gene therapy for driving high levels of expression in both neurons and glial cells (M2a, Fig. 1a). Alternatively, we cloned a 1.4 kb fragment of the *Mecp2* promoter (that following previous studies includes the vast majority of regulatory regions identified so far) with the intent to recapitulate the endogenous *Mecp2* expression pattern (M2b, Fig. 1a). *Mecp2^-/y^* primary neurons were infected with either viruses. As control we used wild-type (WT) neurons infected with a GFP vector (GFP, Fig 1a), and two weeks later were lysed for immunoblotting and genome copy quantification. Surprisingly, using this *Mecp2* promoter fragment, the total viral Mecp2 protein amount remained significantly lower respect to endogenous levels in transduced neurons (Fig. 1b). Conversely, mutant neurons infected with the cassette containing the CBA promoter exhibited a physiological range of total Mecp2 protein. Moreover, similar results were obtained using Mecp2 immunofluorescence staining on infected neuronal cultures (Fig. 1c). This difference was not depended on the relative infection load, since viral copy numbers were equivalent in neurons transduced with either of the two viruses (Fig. 1d). Thus, only the use of the CBA strong promoter was capable of re-establishing Mecp2 protein levels in a physiological range without exceeding viral doses. We, then, asked why and through which mechanism the CBA-*Mecp2* cassette (M2a) could not exceed the endogenous Mecp2 protein range despite the use of a strong promoter and the presence of multiple copies of the virus in the neurons (Fig. 1d). We speculated that the lack of the 5’- and 3’-UTR complete sequences from the *Mecp2* transgene might intrinsically impair protein production. During neuronal development, *Mecp2* transcripts with a long 3’-UTR are highly stabilized leading to progressive Mecp2 protein accumulation^22^. In contrast, alternative *Mecp2* isoforms with shorter 3’-UTRs are less stable and poorly regulated during development. Thus, we sought to compare the relative stability of the viral *Mecp2* transcript respect to the total endogenous *Mecp2* mRNA by measuring its half-life after gene transcription arrest with Actinomycin D (ActD) (Fig. 1e). Using the same qRT-PCR primers and reaction, viral *Mecp2* transcripts were selectively amplified from PHP.eB-*Mecp2* transduced neuronal cultures isolated from *Mecp2* knock-out (KO) embryos, while total endogenous *Mecp2* mRNAs were obtained from WT neuronal cultures (Fig. 1f). Remarkably, viral *Mecp2* transcripts showed significant lower RNA levels respect to the total endogenous *Mecp2* transcripts after 300 min of ActD treatment (51% ± 4%) (Fig. 1f). Thus, the lack of a long 3’-UTR generates an instability-prone *Mecp2* (*iMecp2*) isoform which is significantly destabilized in neuronal cultures. To determine if this reduction in RNA half-life was only determined by the pUTR, we generated an additional cassette with a synthetic assembled 3’-UTR (aUTR) which merged most of the known regulatory elements scattered in the 8,6 Kb long 3’-UTR as previously reported^17^ (M2c, Fig. 1a). In addition, we generated two more vectors, one including the physiological 5’-UTR sequence in combination with the pUTR sequence (M2d, Fig. 1a) and a second lacking the 3’-UTR thus carrying only the polyA (pA) sequence (M2e, Fig. 1a). *Mecp2* RNA levels in neurons transduced with these viral vectors remained very unstable over time compared to endogenous mRNA levels (M2e: 48% ± 13%; M2c: 62% ± 0,2%; M2d 46% ± 20%; M2b 53% ± 3%). Thus, our results confirm that Mecp2 gene expression is regulated by complex mechanisms and multiple factors, especially at RNA level. This complexity can hardly be restricted in the limited capacity of an AAV vector but is essential for the design of future *Mecp2* gene replacement strategies. *Mecp2* mRNA instability was independent by the genotype of the transduced neurons since similar mRNA loss was detected in WT, *Mecp2* mutant and heterozygote neurons (Fig. 1g).

**Figure 1.**
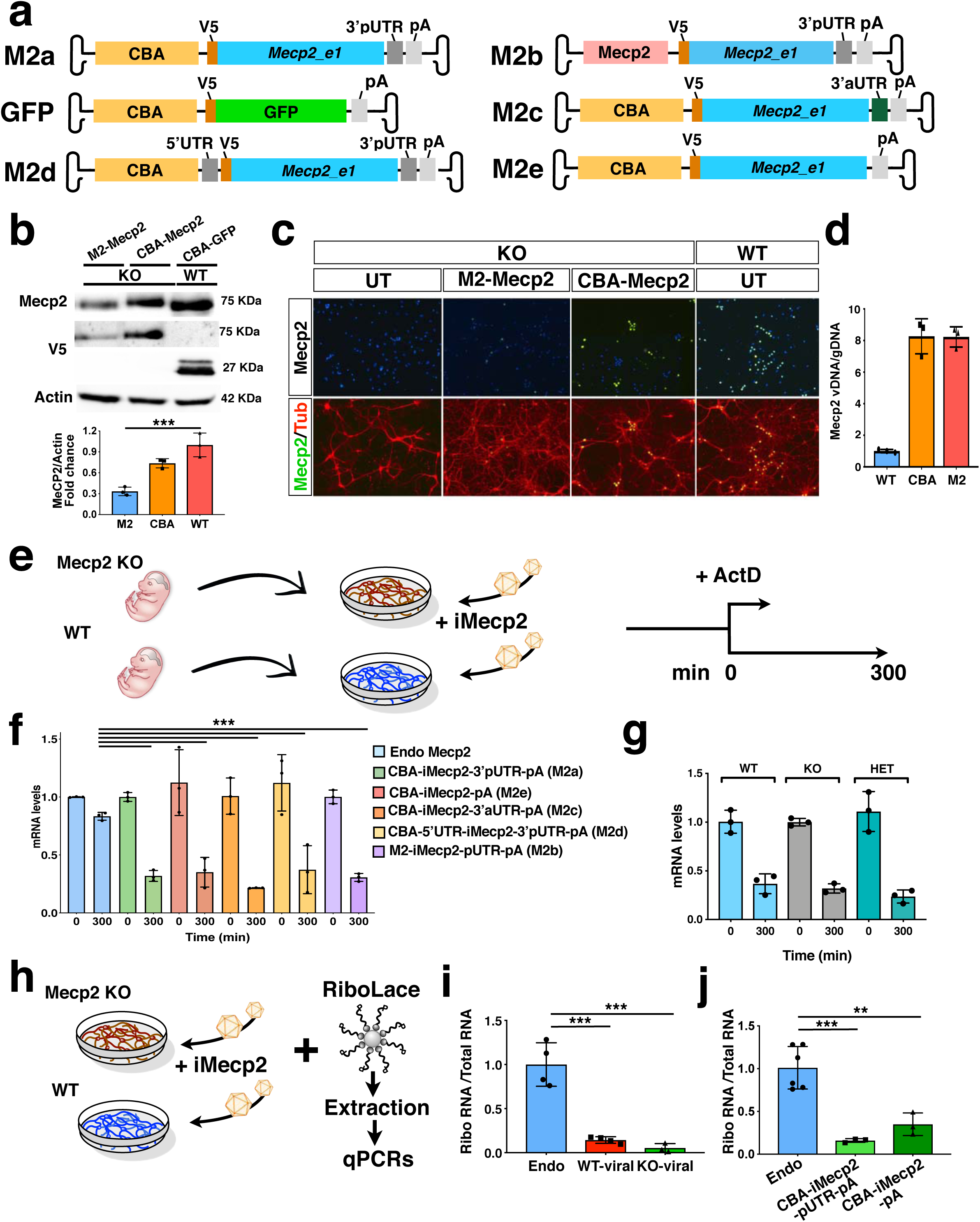
RNA stability and translational efficiency of the viral *Mecp2* transgene. (**a**) Illustration of the AAV vectors expressing V5-tagged *Mecp2_e1* or GFP under the control of Chicken β-Actin (CBA) promoter or *Mecp2* core promoter. Vectors included murine *Mecp2* coding sequence with either its own proximal 3’UTR (3’pUTR, M2a), or a synthetic 3’UTR sequence (3’aUTR, M2c), or both 3’pUTR and 5’UTR (M2d), or no UTR (M2e). (**b**) Western blot analysis for V5, Mecp2 and Actin protein levels in GFP infected (control, CBA-GFP) WT neurons and *Mecp2^-/y^* (KO) neurons infected with CBA-*Mecp2* and M2-*Mecp2*. Quantification was performed using densitometric analysis of Mecp2 relative to Actin signal and expressed in arbitrary units (n = 3) (**c**) Immunostaining of KO (control untreated and infected with M2-*Mecp2* or CBA-*Mecp2*) and wild-type neurons for Mecp2 and TUBB3 (Tub). (**d**) qRT-PCR quantification of viral *Mecp2* DNA copies in Mecp2 KO neurons relative to genomic DNA (n = 3). (**e)** Illustration of the experimental work-flow to study the RNA stability in KO and WT neurons (**f**) RNA stability of endogenous (in KO neurons) and viral (from the 5 different vectors described above in KO neurons) *Mecp2* transcript determined by qRT-PCRs (n = 3, t = 300min were normalized over t=0 values and compared among different treatments). (**g**) RNA stability of viral (CBA-iMecp2-pUTR-pA) *Mecp2* transcript in neurons derived from different genotypes: WT, KO and *Mecp2^+/-^* (Het) determined by qRT-PCRs (n = 3, t = 300min were normalized over t=0 values and compared among different treatments) (**h**) Illustration of the experimental work-flow to study translational efficiency. (**i**) qRT-PCR of viral and endogenous *Mecp2* RNA in the ribosomal fraction normalized on the total RNA in WT and KO neurons (n = 4 endogenous *Mecp2*, n = 4 exogenous *Mecp2* in KO neurons, n = 3 exogenous *Mecp2* in WT neurons). (**j**) qRT-PCR of viral and endogenous *Mecp2* RNA in the ribosomal fraction normalized on the total RNA in WT (n = 6) and KO neurons infected with 2 different viral Mecp2 construct with (CBA-iMecp2-pUTR-pA, n = 3) and without (CBA-iMecp2-pA, n = 3) the 3’-pUTR. Error bars, Standard Deviation (SD). ** p < 0.01, *** p < 0.001, compared groups are indicated by black lines. ANOVA-one way, (b, f, g, i, j) and Tukey’s post hoc test. Scale bar: 100 µm (c).

Next, we sought to determine the translational efficiency of the *Mecp2* viral variant. For this aim, we employed the RiboLace, a novel methodology based on a puromycin-analog which enables the isolation of the ribosomal fraction in active translation with their associated RNAs^23^. Thus, the translational active ribosomal fraction was captured by RiboLace-mediated pull-down from lysates of uninfected WT or PHP.eB-*iMecp2* transduced *Mecp2*-KO neuronal cultures (Fig. 1h). Subsequently, mRNAs were extracted from both the isolated ribosomal fractions and the total lysates and used for RT-qPCR analysis with the same set of *Mecp2* primers. Remarkably, the normalized fraction of viral *Mecp2* transcripts associated with translating ribosomes was reduced by 83% ± 5% compared with that of ribosome-bound endogenous *Mecp2* mRNA (Fig. 1i). A similar reduction was calculated by using a viral *Mecp2* with or without the pUTR suggesting that this impairment is independent by the associated non-coding elements (Fig. 1j). Next, the relative stability of the Mecp2 proteins produced by either the endogenous or the viral gene were assessed in neurons after treatment with cycloheximide (CHX). No difference in Mecp2 protein levels were found up to 8h after CHX administration (Fig. 2a). Finally, we asked whether *MECP2* RNA instability can represent a hurdle also in designing viral cassettes with the human *MECP2* gene. Thus, we employed a pair of male isogenic iPSC lines either control or with a CRISPR/Cas9 induced *MECP2* loss-of-function mutation (Fig. 2b,c). Both iPSC lines were differentiated *in vitro* into cortical neuronal cultures and, then, control and *MECP2* mutant lines were transduced with AAV expressing either GFP or the human version of *iMECP2*, respectively (Fig. 2d,e). Noteworthy, viral *MECP2* transcript stability was severely affected as compared to endogenous RNA levels to an extent comparable to what observed with murine mRNAs (Fig. 2f).

**Figure 2.**
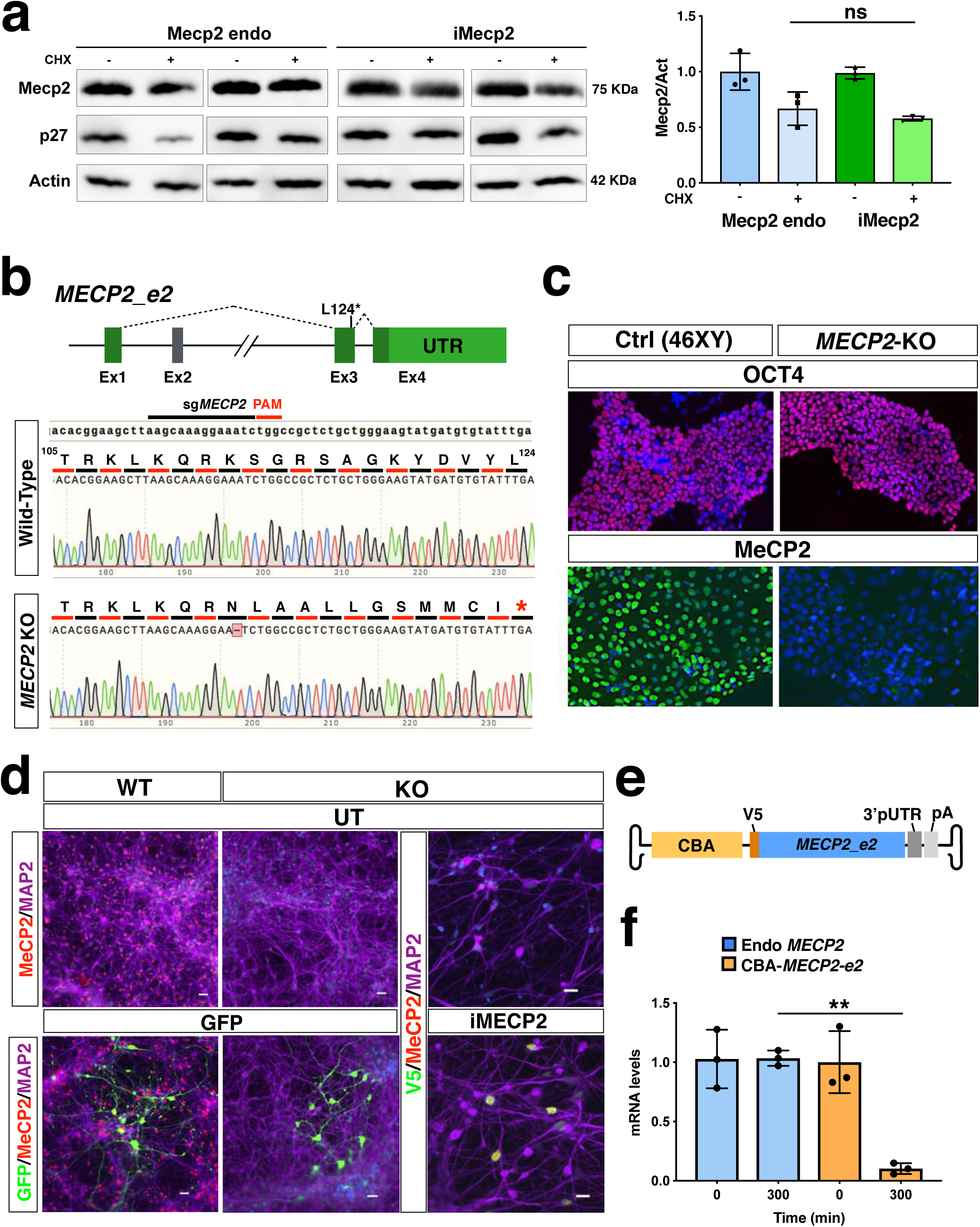
Mouse protein stability and stability of the viral human i*MECP2* construct in iPSC-derived cortical neurons. (**a**) Protein stability was assessed in murine neurons using Western blot analysis to quantify Mecp2 and p27 protein reduction after Cycloheximide treatment (50uM, CHX, 8h) over Actin protein and corresponding densitometric quantification expressed in arbitrary units (right panel) (n = 3). t = 8h were normalized over t = 0 values and compared among different treatments (ns = p > 0.05, ANOVA-one way, Tukey’s post hoc test). (**b**) Illustration of the CRISPR-Cas9 based strategy for genetic editing of male control human iPSCs to obtain *MECP2* KO cells. The sgRNA selected for this approach annealed on exon 3 of the *MECP2* gene and generated a single nucleotide deletion. That resulted in a frameshift of its coding sequence and a premature STOP codon 12 residues downstream. (**c**) Human iPSCs carrying this mutation (*MECP2* KO) maintained their pluripotency marker Oct4 but did not presented detectable MeCP2 protein as tested by immunofluorescence when compared to control cells (Ctrl). (**d**) WT and *MECP2*^-/y^ iPSCs were successfully differentiated into neurons (MAP2 staining) and transduced using PHP.eB vectors carrying either GFP or i*MECP2*. Immunofluorescence respectively for GFP and MeCP2 attested neuronal transduction. (**e**) Schematics of the AAV vector used for RNA stability experiment and expressing V5-tagged *MECP2_e2* under the control of Chicken β-Actin (CBA) promoter. (**f**) *MECP2* RNA stability was determined by qRT-PCRs in WT neurons, to test the endogenous transcript (blue bars, n = 3), and in KO neurons infected with the aforementioned vector, to test the viral transcript (yellow bars n =3). t = 300 min were normalized over t = 0 values and compared among different treatments. UT: untreated.

In summary, these data revealed the crucial role of the post-transcriptional processes in determining the final output of the viral Mecp2 protein levels. Given this limited efficiency in transcript stability and translation efficacy only the use of the CBA strong promoter was effective to sustain Mecp2 protein levels comparable to those found in WT neurons. Since all the viral cassettes caused a similar instability of *Mecp2* transcripts, among them we chose CBA-iMecp2-pUTR-pA (M2a) construct for further *in vivo* studies, and referred to it as *iMecp2*, for instability-prone *Mecp2*.

### Efficient *iMecp2* gene transfer throughout the brain of adult male *Mecp2* mutant mice

To test the efficacy of gene transfer with the *iMecp2* transgene cassette we designed a dose escalation approach to administer 10-fold increasing doses of PHP.eB-*iMecp2* (1×10^9^ vg, 1×10^10^ vg, 1×10^11^ vg and 1×10^12^ vg/mouse) of virus through intravenous delivery in 4 weeks old *Mecp2^-/y^* mice and untreated *Mecp2^-/y^* animals were utilized as controls (Fig. 3a, **Supplementary Fig. 1a**). To determine the exact brain penetration efficiency and neural tissue transduction of the PHP.eB-*iMecp2*, mice for each viral dose were euthanized and brains separated in two halves for immunohistochemistry and immunoblot analysis two weeks after administration. Sections of the infected *Mecp2* mutant brains were stained for Mecp2 or V5 to visualize the viral Mecp2 transduction pattern. The distribution of PHP.eB-*iMecp2* was spread throughout the brain with increasing intensity using higher viral doses (Fig. 3b), as also measured with fluorescence intensity (**Supplementary Fig. 1b**). Quantification of Mecp2^+^ cells respect to DAPI^+^ nuclei in the somatosensory cortex and striatum showed a proportional increase in transduction efficiency from the lowest (1 x 10^9^ vg; 15 ± 3% in cortex; 18 ± 3% in striatum) to the highest dose (1 x 10^12^ vg; 78 ± 3% in cortex; 80 ± 4% in striatum). Brain tissue transduced with 10^11^ vg of PHP.eB-*iMecp2* and immunodecorated for sub-type cellular markers showed that viral transduction was equally efficient in infecting both neuronal and glial cells (**Supplementary Fig. 2a,b**). In particular, the cortical GABAergic interneurons whose dysfunction is a crucial determinant of the RTT phenotype were effectively transduced (**Supplementary Fig. 2a,b**)^24^. Sub-cellular analysis of viral Mecp2 protein distribution confirmed a strong enrichment in nuclear heterochromatic foci, mirroring the genome-wide distribution of endogenous Mecp2 (**Supplementary Fig. 2c**).

**Figure 3.**
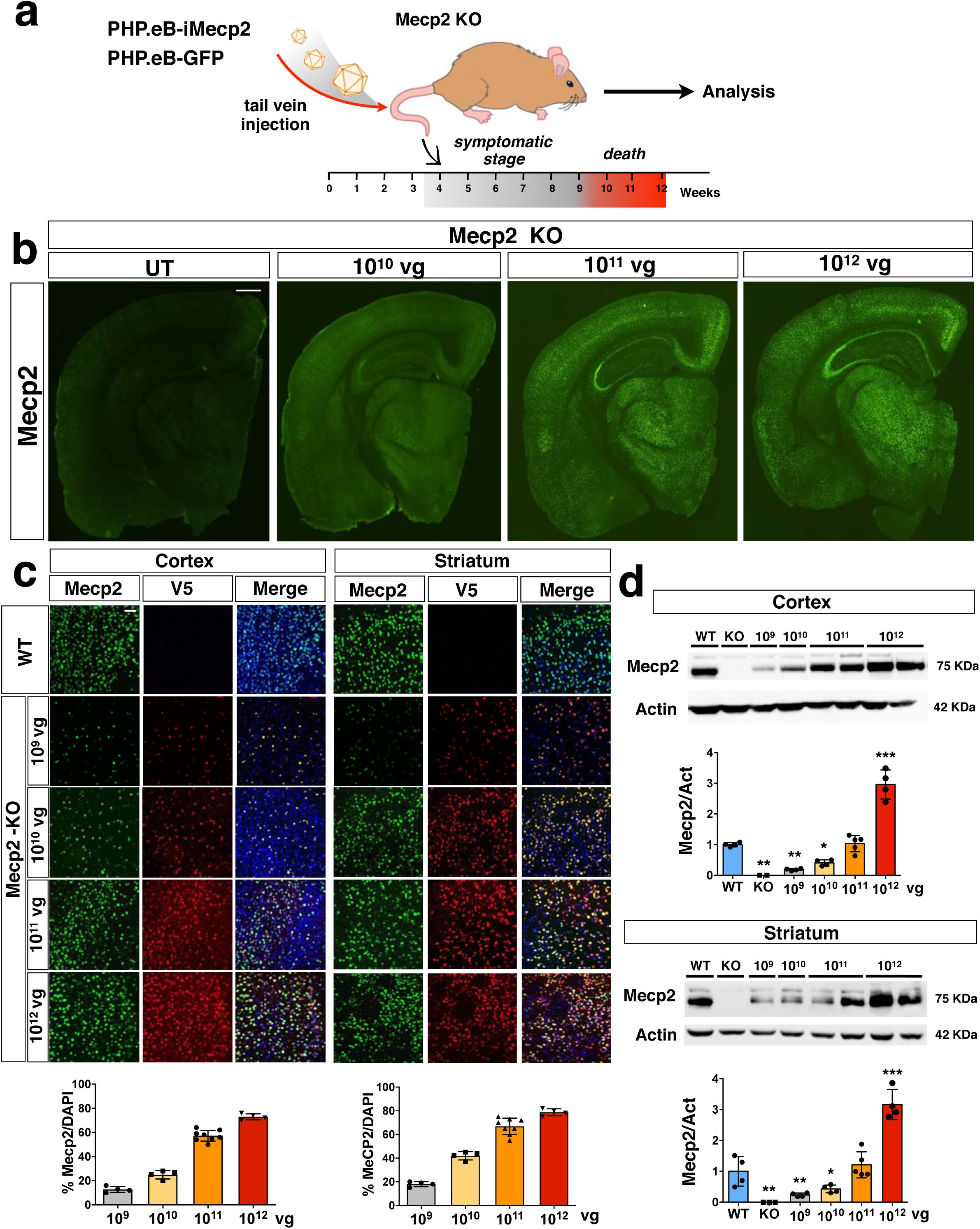
PHP.eB-mediated *iMecp2* gene transfer in symptomatic *Mecp2^-/y^* mouse brains. (**a**) Illustration of the experimental setting to restore the expression of *Mecp2* in symptomatic mutant animals by AAV-PHP.eB systemic transduction. (**b**) Low magnification of Mecp2 immunostaining in brains of KO control untreated (UT) and treated animals (1*10^10^, 1*10^11^, 1*10^12^ vg/mouse). (**c**) High magnification immunostaining for Mecp2 and V5 in cortex and striatum derived from wild-type (WT) and Mecp2 treated KO (1*10^9^, 1*10^10^, 1*10^11^, 1*10^12^ vg/mouse) animals (*Mecp2*-KO). Nuclei were stained with DAPI (merge panels). Bottom panel: bar graphs showing the fraction of Mecp2 positive on the total DAPI positive (n = 4 for 1*10^9^-1*10^10^-1*10^12^; n = 8 1*10^11^ vg/mouse). (**d**) Western blot analysis to quantify Mecp2 over Actin protein levels in cortex (upper panel) and striatum (lower panel) derived from WT, untreated KO and *iMecp2* treated K*O* (1*10^9^, 1*10^10^, 1*10^11^, 1*10^12^ vg/mouse) animals and corresponding densitometric quantification expressed in arbitrary units (n = 4 for 1*10^9^-1*10^10^-1*10^12^; n = 5 1*10^11^ vg/mouse). Error bars, SD. * p < 0.05, ** p < 0.01 and *** p < 0.001 as compared to WT mice (ANOVA-one way with Tukey’s post hoc test). Scale bars: 500 µm (b), 20 µm (c).

Subsequent Western blot analysis revealed increasing total levels of Mecp2 protein in cortical and striatal tissues upon transduction with higher doses of PHP.eB-*iMecp2* (Fig. 3d). In particular, administration of 10^11^ vg of PHP.eB-*iMecp2* resulted in Mecp2 protein levels comparable to those detectable in control brains (Fig. 3d). Conversely, 10^12^ vg of PHP.eB-*iMecp2* triggered a 3-fold increase in *Mecp2* expression respect to endogenous levels (Fig. 3d).

To further assess the efficiency of viral transduction, we measured the number of viral *iMecp2* copies present in the brain and liver of the treated mice. PHP.eB-*iMecp2* treated animals showed a higher number of viral copies in the brain respect to the liver (brain: 15 ± 5, 10^11^ vg; 65 ± 15, 10^12^ vg. liver: 7 ± 2, 10^11^ vg; 55 ± 15, 10^12^ vg) (**Supplementary Fig. 3a**) confirming that the PHP.eB capsid has higher propensity to transduce the neural tissue respect to peripheral organs^20^. In contrast, transgene RNA levels were proportionally less abundant compared to the relative viral genome copy in brain respect to the liver (**Supplementary Fig. 3a**). In addition, despite the significant increase in *iMecp2* genomic copies and total mRNA, protein levels were only marginally augmented in brain and partially in liver (Fig. 3d and **Supplementary Fig. 3b**). This effect was particularly evident in cortical brain samples with the 10^12^ vg dose that compared to wild-type triggered an increase of 80-fold in mRNA, but only of 3-fold in protein. These observations confirmed that *iMecp2* mRNA is poorly transcribed and translated in brain tissue as previously shown in neuronal cell cultures. Altogether, these data clearly highlight the robust efficiency of the PHP.eB capsid to cross the blood-brain barrier in adult mouse brains and to spread throughout the neural tissue transducing large number of cells. Importantly, the four doses of PHP.eB-*iMecp2* which differed by a 10-fold higher titer showed a proportional increase in transduction efficiency in the brain. Hence, this escalation in viral transduction offered a great opportunity to test the extent of phenotypic rescue in *Mecp2* mutant mice depending by the viral gene transfer efficiency and the relative number of cells with restored *Mecp2* expression.

### Severe immune response to exogenous *iMecp2* and its suppression by cyclosporine in transduced *Mecp2^-/y^* mice

Next, we treated mice with increasing doses of PHP.eB-*iMecp2* (1×10^9^ vg, 1×10^10^ vg, 1×10^11^ vg and 1×10^12^ vg) and their control littermates (WT and GFP treated *Mecp2^-/y^* mice, 1×10^11^ vg). Four weeks old *Mecp2^-/y^* mice were intravenously injected and examined over time to monitor the progression of behavioral deficits and the relative efficacy of the treatments. As previously reported, at the age of injection *Mecp2^-/y^* mice in a C57Bl/6 background had already reduced body size and were lighter compared to their WT littermates and between 9-11 weeks of age they experience a sudden weight loss which anticipated the worsening of RTT symptoms and their following decease^4^ (Fig. 4a). Animals were euthanized upon a body weight loss of 20% for ethical reasons. In absence of treatment *Mecp2^-/y^* presented with these features already at 5-6 weeks therefore treatment was never delivered in mice older than this age. A similar lifespan length was observed in mice administrated with two different doses of control PHP.eB-GFP virus (10^11^ vg and 10^12^ vg). Conversely, the *Mecp2^-/y^* mice treated with 10^10^ vg and 10^11^ vg of PHP.eB-*iMecp2* showed a significant increase in lifespan reaching a median survival of 59d and 68d, respectively (Fig. 4a). These beneficial effects were not observed in Mecp2^-/y^ mice injected with the lowest dose of PHP.eB-*iMecp2* (10^9^ vg) whose lifespan remained comparable to that of control treated animals (Fig. 4a). Unexpectedly, all the animals (n = 6) exposed to the 10^12^ vg dose of PHP.eB-*iMecp2* died within two weeks from the viral administration and, then, excluded from the following functional tests (Fig. 4a). These animals did not present with signs of liver toxicity measured as means of blood serum levels of liver proteins and indicated by the absence of alterations in histochemical analysis of liver sections (**Supplementary Table 1**). To determine the general symptomatic stage, the animals were subjected to longitudinal weighting, and a battery of locomotor tests and phenotypic scoring, such as the total grading for inertia, gait, hindlimb clasping, tremor, irregular breathing and poor general conditions that together were presented as the aggregate severity score^7^. *Mecp2^-/y^* mice treated with the median viral doses (10^10^ vg and 10^11^ vg) maintained pronounced locomotor activity and exploratory behavior until few days before their sacrifice (**Supplementary Fig. 4**). In sharp contrast, administration of the lowest dose of virus (10^9^ vg) was not sufficient to exert any detectable beneficial effect (**Supplementary Fig. 4**). Aforementioned immunofluorescence analysis showed a proportional relationship between the increasing doses of virus inoculated in the animals and the enhanced *Mecp2* gene transfer in the brain. With these data we can conclude that reversal of the *Mecp2^-/y^* pathological deficits is related both to transduction efficiency and protein levels achieved in the different districts of the brain. Indeed, when at least 70% of the brain cells were transduced in the cortex and striatum and protein levels were restored up to a physiological range, a sustained behavioral improvement was observed. Similar conclusions were drawn by Robinson et al. (2012) using reactivation of *Mecp2* endogenous gene expression in mice. In contrast brain transduction lower than 15% was not sufficient to attain detectable behavioral improvement.

**Figure 4.**
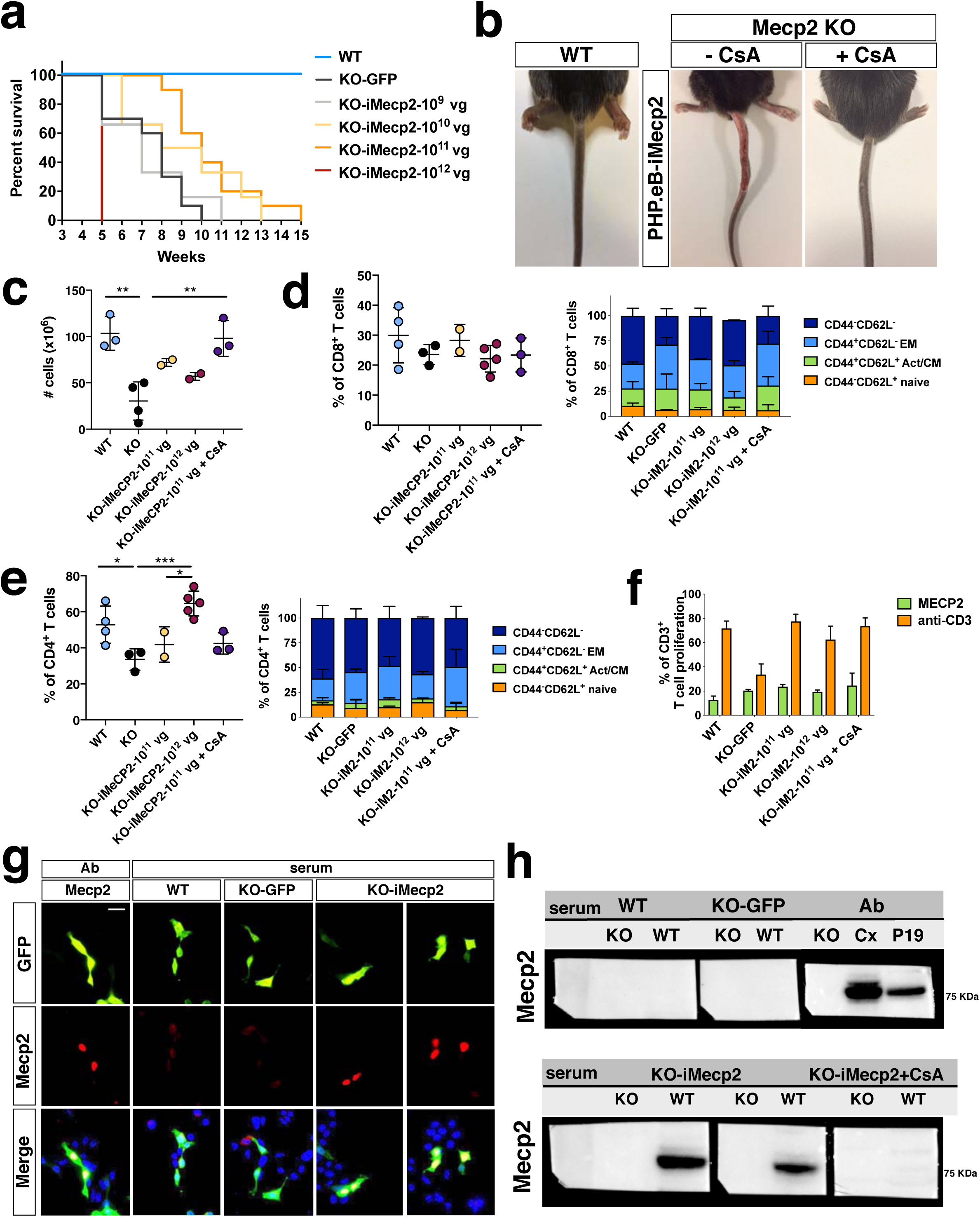
*iMecp2* elicits a strong immune response in *Mecp2^-/y^* but not wild-type mice. (**a**) Kaplan-Meier survival plot for KO mice injected with different doses (1*10^9^ [n = 6], 1*10^10^ [n = 6], 1*10^11^ [n = 10], 1*10^12^ [n = 6] vg/mouse) of PHP.eB-*iMecp2* compared to *KO* treated with PHP.eB-GFP (KO-GFP, n = 10) and WT (n = 14) animals. Mice treated with 1*10^11^ vg/mouse dosage had a median survival period significantly longer than that of vehicle-treated controls (p < 0.01, Mantel-Cox test). (**b**) 9 out of 12 KO mice injected with a 1*10^11^ dose of PHP.eB-*iMecp2* presented with exudative lesions after 2-3 weeks from viral injection (medial panel, representative picture), whereas 7 out of 10 KO mice injected with the same dosage and treated with cyclosporine (CsA) (10 mg/Kg) were robustly improved (right panel, representative picture). (**c-e**) Spleen cells from Mecp2^-/y^ mice left untreated or injected with a 1*10^11^ dose of PHP.eB-*iMecp2* alone or in combination with CsA or with 1*10^12^ dose of PHP.eB-*iMecp2* were counted. Frequencies of CD4+ and CD8+ T cell compartments were quantified in the spleen of treated mice by FACS staining. (**f**) Total splenocytes were labelled with Cell Proliferation Dye eFluor® 670 and stimulated with bone-marrow derived DC transduced with LV-*Mecp2* or with anti-CD3 antibodies and proliferation was measured at day 4 by flow cytometry. Mean ± SEM of *Mecp2^-/y^* mice untreated (n = 3) or injected with a 1*10^11^ dose of PHP.eB-*iMecp2* alone (n=3) or in combination with cyclosporine (n = 3) or with 1*10^12^ dose of PHP.eB-*iMecp2* (n = 2) are shown. (**g**) Detection of immune-specific antibody in sera of KOanimals mock-treated (KO-GFP) or *iMecp2*-treated (KO-*iMecp2*) as well as i*Mecp2*-treated WT animals (WT) was revealed by immunofluorescence assay and compared with a commercial antibody as positive control (Ab). We choose as substrate P19 cells knock-out for the *Mecp2* gene and co-transfected with *GFP* and *Mecp2* expression constructs in order to track with GFP the Mecp2^+^ cells. (**h**) Similar sera were also tested in Western blot analysis using protein extracts form WT and KO tissue respectively as positive and negative controls. WT P19 extract were also used as positive control. Each dot represents one mouse. Error bars, SEM Scale bar: 10 µm. Mann-Whitney U test (two-tailed), with unpaired t-test (c-f), * p < 0.05, ** p < 0.01, compared groups are indicated by black lines.

Nevertheless, independently by the significant behavioral rescue, all the mice treated with 10^11^ vg particles were euthanized because of a severe tail necrosis unrelated to the disease phenotype (Fig. 4b). This phenomenon presented only in 10^11^ vg treated mice (9 out of 12) and occurred 3-4 weeks after AAV injection. It started with a small lesion near the injection site, that progressively expanded to the entire tail and to the body showing a marked hair loss on the lumbar area. These mice were initially unaffected but while necrosis aggravated weight and general health started to be compromised and sacrifice was necessary. Overall none of the mice treated with lower doses of *iMecp2* or GFP virus presented with this adverse symptom.

Given the systemic delivery of the virus in Mecp2 deficient mice, we set out to assess whether the immune response to the transgene could explain the health deterioration observed in the treated mice. This was unanticipated at least since none of the previous attempts of systemic gene therapy have considered or described such response to happen^15,16,21^. However, Mecp2 has been implicated in the regulation of immunity^25^ and of FoxP3 expression, a transcription factor required for the generation of regulatory T (Treg) cells, during inflammation^26^. Thus, we hypothesized that the lack of Treg-mediated regulation was responsible for the strong immune response observed in *Mecp2^-/y^* mice exposed to PHP.eB-*iMecp2*. Indeed, treatment with cyclosporine A (CsA) of *Mecp2^-/y^* mice exposed to a 10^11^ vg dose of PHP.eB-*iMecp2* resulted in a striking amelioration of the general health conditions (Fig. 4b). The analysis of the frequency of cells in the spleen of *Mecp2^-/y^* mice exposed to 10^11^ vg and 10^12^ vg of PHP.eB-*iMecp2* (KO-*iMecp2*-10^11^ vg and KO-*iMecp2*-10^12^ vg, respectively) showed the increased number of splenocytes harvested from the latter mice compare to untreated *Mecp2^-/y^* controls (Fig. 4c,d). Interestingly, *Mecp2^-/y^* mice showed a significantly lower number of splenocyte compared to WT littermates, in line with the general status of inflammation due to spontaneous activation of T cells in these mice^26^. No major differences were observed in CD8^+^ T cell compartments in *Mecp2^-/y^* mice untreated or exposed to 10^11^ vg and 10^12^ vg of PHP.eB-*iMecp2* virus (Fig. 4d). Interestingly, *Mecp2^-/y^* mice exposed to the higher PHP.eB-*iMecp2* dose (10^12^ vg), analyzed two weeks post treatment, showed a significantly higher frequency of CD4^+^ T cells compared to untreated control mice (Fig. 4e). Hence, we can speculate that treatment with the higher dose of PHP.eB-*iMecp2* virus in *Mecp2^-/y^* mice lacking Treg-mediated regulation led to an uncontrolled inflammatory response associated to the expansion of CD4^+^ T cells that resulted in rapid death. We, next, investigated the induction of Mecp2-specific immune response by analyzing proliferation of T cells in response to Mecp2, cytotoxic activity of CD8^+^ T cells, and anti-Mecp2 antibody production in *Mecp2^-/y^* mice exposed to PHP.eB-*iMecp2* virus. Neither Mecp2-specific T cells nor Mecp2-specific cytotoxic CD8+ T cells were detected in *Mecp2^-/y^* mice exposed to PHP.eB-*iMecp2* virus (Fig. 4f, and data not shown). However, an increased non-specific (anti-CD3-stimulated cells) proliferative response was observed in *Mecp2^-/y^* mice exposed to PHP.eB-*iMecp2* virus compared to untreated mice (Fig. 4f). Finally, we detected anti-Mecp2 antibodies in the sera of *Mecp2^-/y^* mice exposed to PHP.eB-*iMecp2* virus but not in WT mice exposed to PHP.eB-*iMecp2* virus or in control untreated *Mecp2^-/y^* mice (Fig. 4g,h). Importantly, treatment with CsA prevented the induction of anti-Mecp2 antibodies (Fig. 4h). Overall, these studies indicate, for the first time, the induction of uncontrolled proliferation of T cells and induction of Mecp2-specific antibodies in *Mecp2^-/y^* mice exposed to PHP.eB-*iMecp2* virus, which can be overcome by immunosuppression. This severe immune response to the transgene can explain the rapid phenotypic deterioration and premature death occurred to the *Mecp2^-/y^* mice treated with the highest dose of the therapeutic virus (10^12^ vg). This conclusion is corroborated by the fact that WT and females *Mecp2^+/-^* animals exposed to the same dose virus at a comparable dose did not develop any of these complications (see below).

### PHP.eB-*iMecp2* and Csa co-treatment triggers a significant behavioral rescue of male *Mecp2* mutant mice

Next, *Mecp2^-/y^* mice were co-treated with 10^11^ vg PHP.eB-*iMecp2* or PHP.eB-GFP with daily injections of Csa starting 1 day before viral administration and longitudinally profiled for survival rate, general healthy conditions and motor behavior. Remarkably, CsA-treated respect to untreated *Mecp2* deficient mice administrated with PHP.eB-*iMecp2* showed a triplicated mean survival rate, reaching up to 38 weeks of age (Fig. 5a; median survival: KO-GFP: 52d; KO-GFP +Csa: 56d; KO-iMecp2-10^11^: 68d; KO-iMecp2-10^11^ + CsA: 168d). Unfortunately, not all iMecp2-treated animals responded to CsA. Uncomplete immunosuppression lead to tail lesion and premature sacrifice in 3 out of 10 cases, nonetheless all mice presented significant behavioral improvement. In fact, this group of mice exhibited only very low grades (< 3) in the aggregate severity score for an extensive period of time reflecting their general good healthy conditions (Fig. 5b, **Supplementary movie**). In particular, recovery in inertia, gait, hindlimb clasping, and irregular breathing were fully maintained over long time. Only some degree of tremors was still manifested in some of these mice accounting for the scores recorded in this test. This manifestation is likely associated with abnormalities in the mutant peripheral nervous system which is known to be not efficiently transduced by this particular AAV capsid^20^. Motor behavior was strongly rescued after gene therapy as shown by a significant recovery in the rotarod performance (Fig. 5c) and spontaneous locomotor activity (Fig. 5d) up to 15 weeks. *Mecp2^-/y^* mice were previously reported to have reduced anxiety levels^27^. In fact, control treated mutant mice tarried and moved less within the elevated closed arms, while PHP.eB-*iMecp2* treated mice displayed anxiety levels more similar to WT animals (Fig. 5e). Finally, sustained improvement in motor coordination and balance was also evident in the beam balance test (Fig. 5f,g). In summary, this phenotyping assessment showed how a single injection of PHP.eB-*iMecp2* adjuvated by the immunosuppressing treatment can elicit a robust and long-lasting recovery in survival (up to 38 weeks). More importantly the general health state and behavioral skills in *Mecp2^-/y^* mice was also restored to levels comparable to wild-type animals. This is the first clear evidence that a gene therapy approach could rescue for a long period RTT mice and significantly ameliorate their general condition and motor behaviour. The association of a significant life-span increment and evident phenotypic amelioration is the main goal of gene therapies applied to life treating diseases.

**Figure 5.**
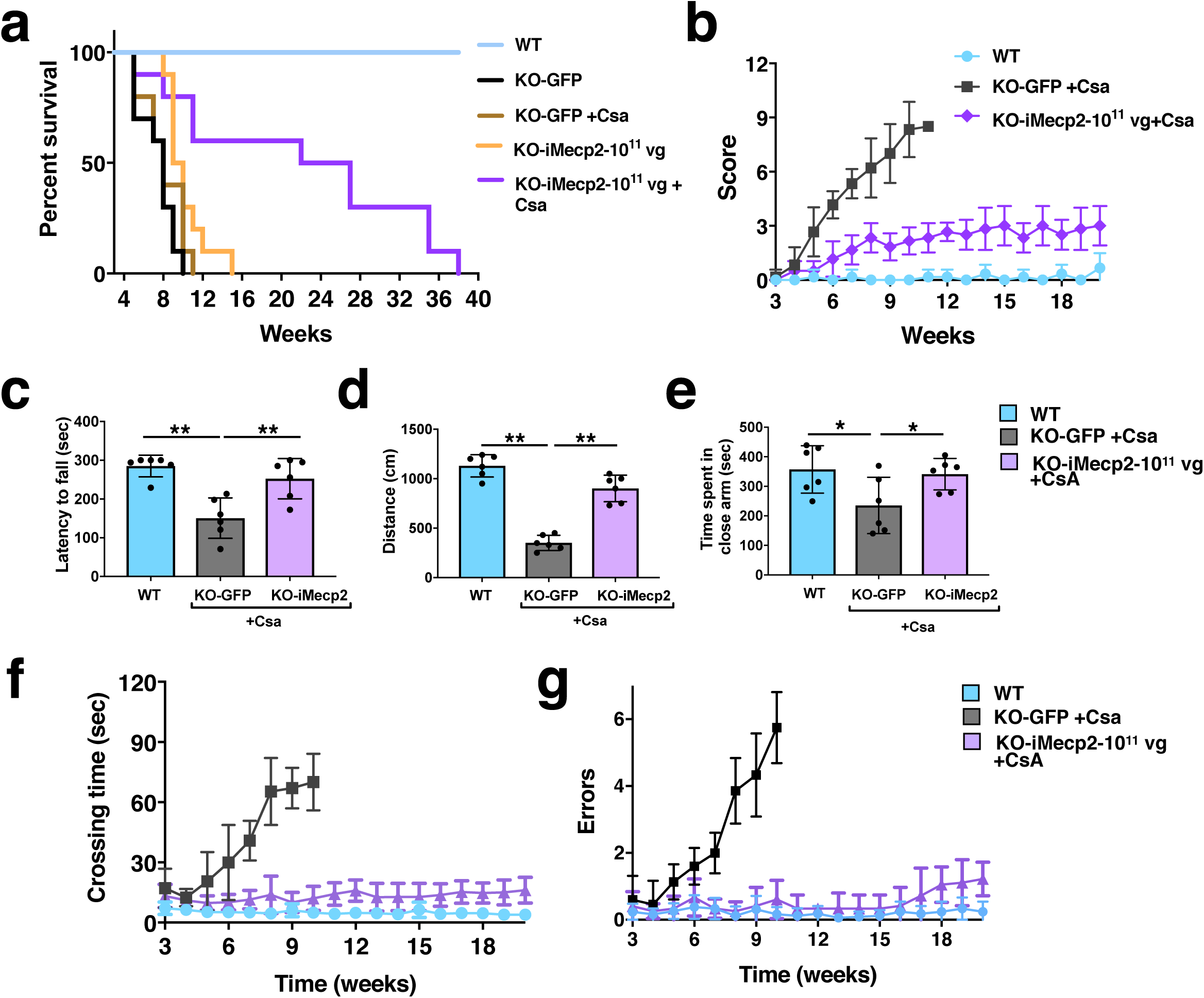
Behavioral rescue of symptomatic *Mecp2^-/y^* animals after PHP.eB-*iMecp2* treatment in combination with cyclosporine. (**a**) Survival curve of KO mice GFP-treated alone (n = 10, black line) or in combination with cyclosporine (CsA) (n = 6, brown line) or injected with a 1*10^11^ dose of PHP.eB-*iMecp2* alone (n = 10, orange line) or in combination with cyclosporine (CsA) (n = 10, violet line). As control, WT littermates were used (n = 12, blue line). (**b**) General phenotypic assessment evaluated through the aggregate severity score (p < 0.05 versus KO-GFP + Csa in 1*10^11^ + Csa [5^th^-11^th^ wk]). (**c-e**) Evaluation of the latency to fall from the Rotarod analysis (**c**), the distance of the spontaneous locomotor activity in a field arena (**d**) and the time spent in close arm in the elevated plus maze (**e**) 5 weeks after treatment and control mice (** p < 0.01 and *** p < 0.001 as compared to WT mice and KO-*iMecp2* + Csa; n = 6 mice per groups). (**f-g**) Evaluation of motor coordination through beam balance test quantified as crossing time (f, p < 0.05 versus KO-GFP + Csa in 1*10^11^ + Csa [6^th^-10^th^ wk]) and number of errors (g, p < 0.05 versus KO-GFP + Csa in 1*10^11^ + Csa [7^th^-10^th^ wk]). Error bars, SD. ANOVA-two way (b, f, g) or ANOVA-one way (c, d, e) with Tukey’s post hoc test.

### PHP.eB-*iMecp2* improves disease symptoms in female *Mecp2* heterozygous mice

*Mecp2^-/y^* mice are an extremely severe model of RTT but does not reflect the mosaic gene inactivation occurring in girls with RTT. Moreover, the strong immune response has complicated the readout of the symptomatic recovery of gene therapy in these animals. Thus, we thought to validate our approach in female *Mecp2^+/-^* mutant mice. To this end, 5 months old *Mecp2^+/-^* animals were intravenously injected with 10^11^ vg of either PHP.eB-*iMecp2* or control virus and examined over time up to 11 months post-treatment (**Supplementary Fig. 5a**). For these experiments, we selected only a single viral dose that sustained a general brain transduction rate between 65% and 90% and raising Mecp2 protein levels similar to those in WT (**Supplementary Fig. 6**). As previously reported, *Mecp2^+/-^* females started to exhibit pathological signs from 10 months of age acquiring breathing irregularities, ungroomed coat, inertia and hindlimb clasping^4^. While control treated mutant females showed a pronounced and sustained weight gain over time, the animals with the viral therapy gradually normalized their weight reaching values similar to those of unaffected mice at 15 months (**Supplementary Fig. 5b**). In the severity score the control treated *Mecp2^+/-^* females progressed to values over 4, whereas animals given the therapeutic virus rarely overcome a score beyond 2 in the entire observation period (**Supplementary Fig. 5c**). Total mobility assessment in the open field showed that at 13 months old PHP.eB-*iMecp2* treated mice have a significant increase in the travelled distance and general activity respect to the control treated group matching the general performance of WT females (**Supplementary Fig. 5d**). Likewise, in the beam balance test the therapy largely preserved the motor skills of the *Mecp2^+/-^* mice that, in contrast, were partially lost in control treated mutant mice (**Supplementary Fig. 5e,f**). Collectively, these observations provide evidence that the PHP.eB-*iMecp2* treatment sustained a significant and long-term protection from symptomatic deterioration improving the health conditions and reducing the locomotor phenotype in female *Mecp2^+/-^* mice.

### PHP.eB-*iMecp2* reduces molecular and gene expression alterations in treated *Mecp2^-/y^* mice

In order to examine the molecular alterations downstream to Mecp2 loss and evaluate whether gene therapy with PHP.eB-*iMecp2* might sustain any discernable recovery, we performed global gene expression analysis by RNA-Seq of whole cerebral cortical tissue from 9 weeks old *Mecp2^-/y^* mice inoculated with either 10^11^ vg of PHP.eB-*iMecp2* or control virus and WT littermates. Computational analysis identified 1876 differential expressed genes (DEGs) with p < 0.05 significance between *Mecp2^-/y^* and control mice roughly divided in two equal groups between up- and down-regulated genes in mutant mice (Fig. 6a). This dataset significantly overlaps with the results published in a previous RNA-seq study which profiled the same brain tissue at a comparable mouse age, confirming the consistency of our results (data not shown)^28^. The number of significant DEGs between Mecp2^-/y^ mice administrated with therapeutic or control virus was 1271 with a small increase in upregulated genes. However, only a third of DEGs were shared between viral transduced and untreated *Mecp2^-/y^* mice, while the remaining DEGs of the mutant mice were normalized in the treated counterparts (Fig. 6b,d,e). This data suggested that the viral treatment was able to correct a large fraction of gene expression changes associated with the RTT phenotype. However, a large set of DEGs was only associated with the treated *Mecp2^-/y^* mice and, thus, to uncover their significance we performed Gene Ontology functional enrichment analysis (GO). Remarkably, most of these DEGs were associated with immunological pathways such as immune response, immune system regulation and inflammation (Fig. 6c). Hence, these results corroborated at the molecular level the previous observations on the strong immune response mounted in the treated *Mecp2^-/y^* mice against the inoculated transgene. We, then, performed gene ontology analysis on the DEG dataset enriched in the *Mecp2^-/y^* mice but normalized after gene therapy. Interestingly, the most significant enrichment was in metabolic networks associated with lipid biosynthesis/transport and in particular cholesterol metabolism (Fig. 6f). Previous studies have shown brain and peripheral cholesterol levels are altered in *Mecp2* mutant mice and patients’ specimens^29–31^. In addition, gene responsible for cholesterol biosynthesis has been shown to be downregulated in severe symptomatic *Mecp2^-/y^* animals^29^. Remarkably, a large component of the molecular pathway for cholesterol production was downregulated in *Mecp2^-/y^* mice, but significantly rescued in PHP.eB-*iMecp2* transduced animals (Fig. 6g). RT-qPCRs on independent cortical tissue lysates confirmed that gene expression levels of crucial enzymes in the cholesterol biosynthesis such as Squalene epoxidase (*Sqle*), NAD(P)-dependent steroid deydrogenease-like (*Nsdhl*) and Methylsterol monooxygenase 1 (*Msmo1*) were significantly restored by PHP.eB-*iMecp2* gene therapy (Fig. 6i). An additional molecular group highly divergent between transduced and control *Mecp2^-/y^* mice included genes encoding for potassium (Kv) channels. This class of ion channels serve diverse functions including regulating neuronal excitability, action potential waveform, cell volume and fluid and pH balance regulation. Interestingly, it was previously reported that *Mecp2* deficiency leads to decreased *Kcnj10*/*Kir4.1* mRNA levels and related currents in mutant astrocytes^32^. We confirmed reduced *Kcnj10* transcripts together with gene deregulation of other Kv channels in *Mecp2^-/y^* mice (Fig. 6h,i). Among others, the potassium channel gene *Kcnc3*, associated with ataxia and cognitive delay in humans, was downregulated in *Mecp2* mutants and normalized after the PHP.eB-*iMecp2* treatment (Fig. 6h,i). We previously showed that *Mecp2* mutant brains exhibit reduced mTOR signaling with diminished phosphorylation of phospho-S6 (pS6)^33^. Remarkably, overall levels of pS6 on Ser234/235 were significantly increased in brain tissue transduced with the PHP.eB-*iMecp2* virus (Fig. 6j). In summary, PHP.eB-*iMecp2* gene therapy sustained a wide recovery of the abnormal gene expression in the *Mecp2* mutant brain tissue and elicited the rescue of the global impairment affecting transcriptional and translational processes upon *Mecp2* gene loss.

**Figure 6.**
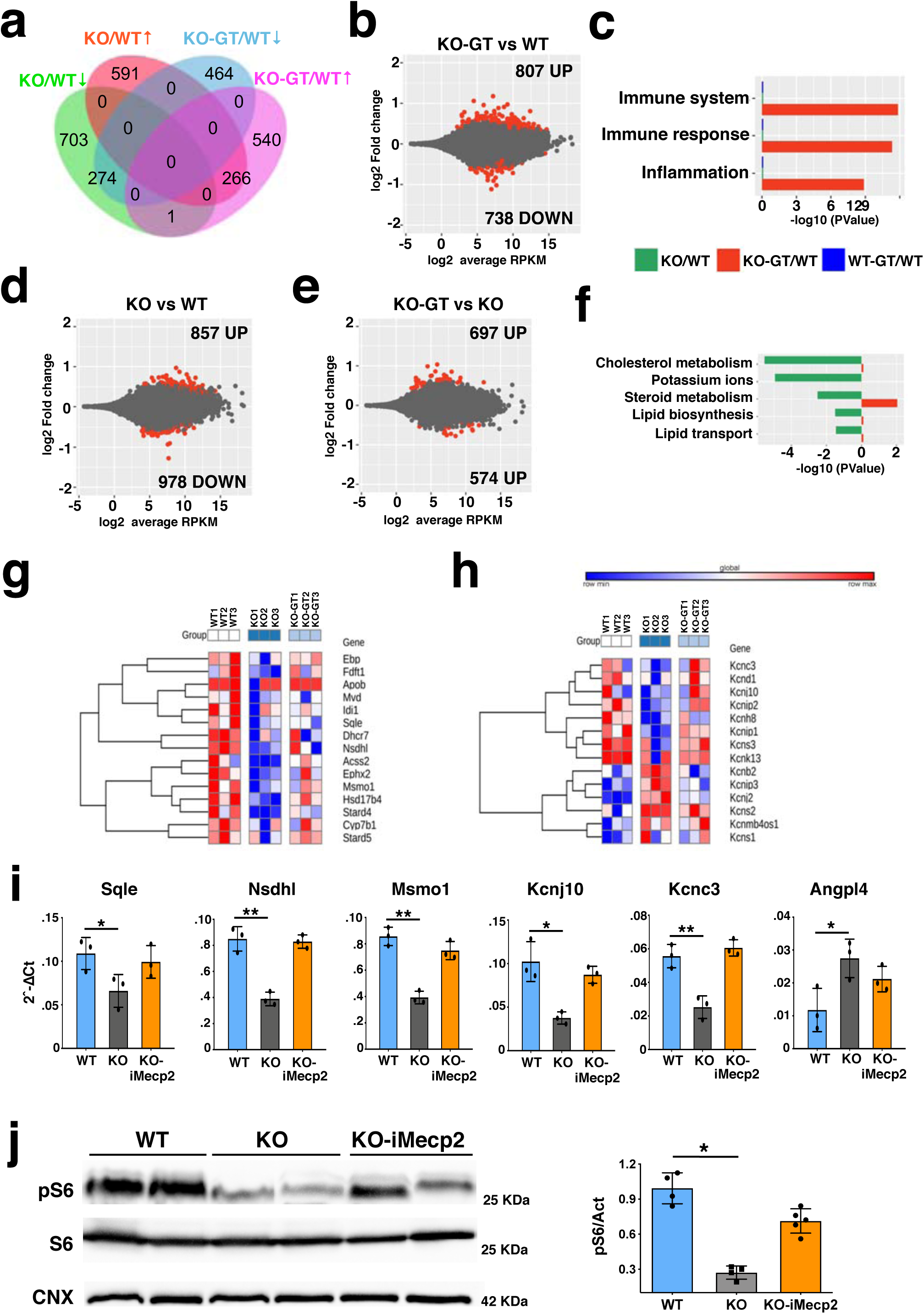
Global gene expression profile of *Mecp2^-/y^* cortices transduced with PHP.eB-*iMecp2*. **(a)** Venn diagram showing the genes differentially expressed in the two comparisons, namely *Mecp2* KO (supplied with a mock treatment with PHP.eB-GFP 10^11^ vg/mouse, n = 3) vs WT (n = 3), and *Mecp2* KO-GT (after gene therapy treatment, PHP.eB-*iMecp2* 10^11^ vg/mouse, n = 3) vs WT. **(b)** MA plot showing gene expression fold changes as a function of the average gene expression in the *Mecp2* KO-GT vs WT comparison. (**c**) Representative gene ontology categories highlighting the enrichment for immune response-related datasets being overrepresented in the *Mecp2* KO-GT vs WT comparison. (**d**) MA plot showing gene expression fold changes as a function of the average gene expression in the *Mecp2* KO vs WT comparison. (**e**) MA plot showing gene expression fold changes as a function of the average gene expression in the *Mecp2* KO-GT vs KO comparison. (**f**) Representative gene ontology categories highlighting the enrichment for lipid metabolism-related datasets being overrepresented in the *Mecp2* KO vs WT comparison, but not in the *Mecp2* KO-GT vs KO comparison. Heatmap showing relative expression of genes belonging to the lipid metabolism-related pathways (**g**) or Potassium ion transmembrane transport group (**h**). (**i**) RT-qPCRs of selected transcript of interest such as *Sqle Nsdhl*, *Msmo1*, *Kcnc3* and *Angpl4* being downregulated in *Mecp2* KO and rescued after gene therapy. Red dots in **b**, **d**, and **e** depict differentially expressed genes with FDR ≤ 0.1. (**j**) Representative Western blot and quantitative analysis for the ribosomal protein S6, its phosphorylated form (pS6) and a normalizer (CNX) from cortical tissues of wild-mice and *Mecp2* KO transduced with GFP or *iMecp2* vector. Error bars, SD. * p < 0.05, ** p < 0.01 as compared to WT mice (ANOVA-one way with Tukey’s post hoc test).

### Systemic gene transfer of *iMecp2* is not detrimental for wild-type mice

Systemic delivery of PHP.eB-*iMecp2* exerted a large symptomatic reversibility both in male and female Mecp2 mutant mice. To further extend these observations and validate the safety of this treatment we decided to administer the same treatment to WT C57BL/6J adult mice. Animals were administrated with either 10^11^ vg or 10^12^ vg of PHP.eB-*iMecp2* or left untreated (n = 9 each) and closely inspected over time. Next, two animals per group were euthanized 3 weeks after viral inoculation and brain processed for histological analysis. *iMecp2* gene transfer efficiency was evaluated by V5 immunofluorescence which distinguished the viral from the endogenous *Mecp2*. According to aforementioned results, brain transduction efficiency was very high with a net increase of 20% between the lower and the higher viral dose (cortex: 45% ± 8% 10^11^ vg, 68% ± 7% 10^12^ vg; striatum: 58% ± 7%, 10^11^ vg; 82% ± 5%, 10^12^ vg) (Fig. 7a,b). Viral copy number analysis confirmed a significant and prevalent targeting of the virus in the neural tissues respect to the liver (**Supplementary Fig. 7**). Despite the high expression of the transgene, the total Mecp2 protein levels in cortex and striatum were only increased by 30% and 60% upon transduction with 10^11^ vg and 10^12^ vg of virus, respectively as assessed by immunoblotting and immunofluorescence intensity (Fig. 7c, **Supplementary Fig. 8**). General health state and behavior were, then, scored in the remaining treated animals up to 12 weeks after treatment. During this time, growth curve, locomotor activity and fine coordination were slightly different between transduced and control animals (Fig. 7d,e). General health state examination (severity score) revealed some breathing irregularities and tremor at rest which slightly increased the scoring output in the treated animals although with minimal difference compared to WT animals (Fig. 7f). Altogether, these observations indicated that high doses of PHP.eB-*iMecp2* virus did not exert deleterious effects in WT animals in this window of time. Importantly, even the 10^12^ viral dose, which is 10-fold higher than the amount used in *Mecp2^-/y^* mice to trigger substantial beneficial effects, was incapable to trigger a consistent deleterious outcome in the mice except for mild alterations. Next, we performed global gene-expression analysis by RNA-seq from cerebral cortical tissues of animals untreated or inoculated with a 10^11^ vg dose. Remarkably, bioinformatics analysis did not distinguish genes with significantly different expression between the two conditions (p < 0.05) (Fig. 7g,h). Collectively, widespread brain transduction of PHP.eB-*iMecp2* in WT animals elicited only a minimal increase in total Mecp2 levels which was not sufficient to exert neither significant behavioral symptoms nor abnormal gene expression changes.

**Figure 7.**
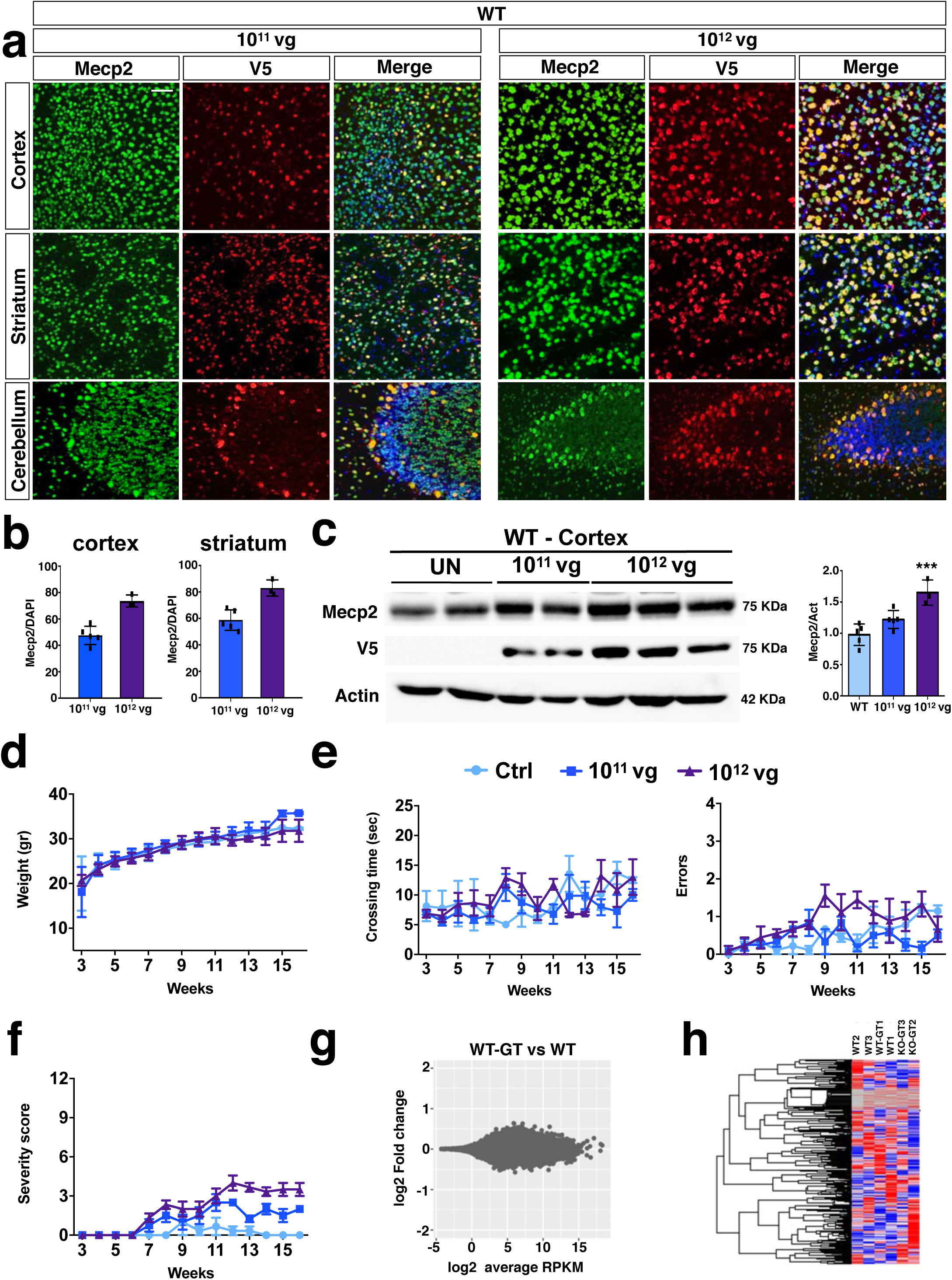
PHP.eB-mediated *iMecp2* gene transfer in wild-type mice is unharmful. (**a**) High magnification immunostaining for Mecp2 and V5 in cortex, striatum and cerebellum derived from WT brains treated with PHP.eB*-iMecp2* (1*10^11^ and 1*10^12^ vg/mouse). Nuclei were stained with DAPI (merge panels). (**b**) Bar graphs showing the fraction of V5 positive on the total DAPI positive in cortex and striatum (n = 6 for 1*10^11^ and n = 3 for 1*10^12^ vg/mouse). (**c**) Western blot analysis to quantify Mecp2 over Actin protein levels in cortex (left panel) and striatum (right panel) derived from WT, untreated and treated with *iMecp2* (1*10^11^ and 1*10^12^ vg/mouse) animals and corresponding densitometric quantification expressed in arbitrary units (right panel) (n = 6 for WT, n = 6 for 10^11^ and n = 3 for 10^12^ vg/mouse) *** p < 0.001 as compare to wild-type (WT) untransduced mice (ANOVA-one way, Tukey’s post hoc test). Twice a week, mice were tested for (**d**) body weight, (**e**) beam balance test and quantified as crossing time (left) and number of errors (right) and (**f**) general phenotypic assessment evaluated through aggregate severity score (WT untreated [n = 8] and treated with *iMecp2* virus 1*10^11^ [n = 6], 1*10^12^ [n = 6], vg/mouse). (**g**) MA plot showing gene expression fold changes as a function of the average gene expression in the *Mecp2* WT-GT (after gene therapy treatment, PHP.eB-*iMecp2* 10^11^ vg/mouse, n = 3) vs WT (n=3) comparison (**h**) Red dots depict differentially expressed genes with FDR ≤ 0.1. Heatmap showing relative expression of 1000 random genes in WT and WT-GT, highlighting the lack of differentially expressed genes in the two groups. Error bars, SD. Scale bar: 50 µm.

## Discussion

Herein, we provided solid evidence that the global brain transduction of the *iMecp2* transgene by PHP.eB-mediated delivery was capable to significantly protect male and female *Mecp2* mutant mice from the symptomatic hallmarks of the RTT phenotype. Importantly, an initial set of viral infections on primary neuronal cultures enabled us to select the most suitable configuration of the transgene cassette in order to achieve Mecp2 protein levels within a physiological range. Our results showed that the lack of a long 3’-UTR sequence destabilizes *Mecp2* mRNA significantly reducing its relative half-time. Moreover, all the combinations of *Mecp2* regulatory regions tested in this work did not enhance the *iMecp2* transgene RNA stability. Thus, more work is necessary to identify additional *Mecp2* transcript stabilizing elements in the long 3’UTR. Additionally, using the RiboLace system to determine the amount of *Mecp2* transcripts associated with active translating ribosomes, our data showed a major loss in translation efficiency of the viral compared to the endogenous mRNA. These results strongly imply the strong functional impact of untranslated elements in the post-transcriptional control of Mecp2 expression. Along these lines, it was previously reported that endogenous Mecp2 protein levels correlate poorly with mRNA levels during development and in different adult tissues^34^. For this reason, we favored to combine the *Mecp2* cDNA with the CBA strong promoter, building the *iMecp2* cassette, in order to reach physiological range of Mecp2 proteins levels. Based on our *in vitro* results we be believe that more broadly, a preliminary *in vitro* screening in primary neuronal cultures is a valuable tool to characterize the properties of newly designed transgene cassette and vector transduction.

We decided to package the *iMecp2* construct in the PHP.eB that was selected for its unprecedented capability to efficiently penetrate the brain after intravenous delivery and widely transduce neuronal and glial cells^20^.

We showed that administration of high dose of PHP.eB-*iMecp2* at the initial symptomatic stage can ameliorate disease progression with robust beneficial effects and significant lifespan extension. Moreover, we observed that both transduction efficiency and Mecp2 protein levels can impact the final outcome of phenotypic recovery. Indeed, animals treated with 10^9^ vg dose of PHP.eB-*iMecp2* (which transduced less than 15% of brain cells cortex and striatum) did not present significant therapeutic effects. The intermediate doses (10^10^ e 10^11^) had respectively moderate and robust effect in reverting phenotypic symptoms. However, in this last case the beneficial effects of the treatment were concealed by an adverse effect consisting of a strong immune response against the transgene. Importantly, this adverse complication is the result of using the male *Mecp2* KO mice that have no Mecp2 from birth and therefore recognize the therapeutic gene product as *nonself*. This is not the case for human female patients that are a mosaic of mutant and WT *Mecp2* cells, moreover even male hemizygous patient rarely present with protein complete loss, thus immune response against this transgene is unlikely to occur in human patients. Nonetheless this is a clinical aspect that need to be carefully considered during patient selection. It was initially surprising to us why former studies of *Mecp2* gene therapy in KO mouse strains did not highlight such adverse event^15,16^. A possible explanation could be related to the different tropism of the PHP.eB in comparison to AAV9.

The life-span of animals treated with 10^12^ vg/mouse dose was not long enough to justify a similar response thus indicating a different mechanism of toxicity. This serious inflammatory response to the transgene, associated to the expansion of CD4^+^ T cells, can explain the sudden death of the *Mecp2^-/y^* animals treated with the highest PHP.eB-*iMecp2* dose (10^12^ vg). In fact, a similar dose of the therapeutic virus in WT animals was free of deleterious effects. Despite previous reports^15^, we could not detect any sign of toxicity in livers treated with PHP.eB, irrespectively of the transgene or the mice genotype, thus ruling out epatotoxicity in these conditions.

Beyond this immune reaction, we could not score any additional adverse manifestations directly caused by the therapy neither in mutant *Mecp2* or WT animals. This is consistent with the mild increase in total Mecp2 levels achieved by the *iMecp2* transgene both in a mutant or WT background. Nonetheless we cannot exclude that the deleterious effect observed in mice treated with the highest dose (10^12^ vg/mouse) could be related to an increase of Mecp2 above physiological level (∼3 fold). In fact, abrupt *Mecp2* reactivation in Mecp2 knock-out mice has been reported to often lethal^7^.

Nevertheless, viral (10^11^ vg) and CsA co-treatment was able to promote an unprecedented life-span recovery and robust improvement in general health state, locomotor activity and coordination. After months of cyclosporine treatment, the mice showed a deterioration in quality life and eventually had to be euthanized mainly for symptoms related to immunosuppression or relapsing immune response. PHP.eB-*iMecp2* intervention significantly corrected also the abnormal gene expression alterations observed in *Mecp2^-/y^* mouse cortical tissue, including the reduced expression in multiple molecular components of the cholesterol pathway and some genes encoding for Kv channels. More broadly, PHP.eB-*iMecp2* gene therapy sustained a strong recovery of the genome-wide transcriptional and mTOR-mediated translational processes affected in *Mecp2* deficient mice.

The molecular characterization *Mecp2^-/y^* mice rescue combined with its phenotypic analysis is a powerful tool to assess molecular pathways involved in the disease and more importantly in its recovery.

Taken together, the diffuse penetration of the PHP.eB in the adult mouse brain parenchyma is a unique property among all the recombinant viral strains in current use. Recently, two studies concomitantly discovered Ly6a as the endothelial receptor that allows PHP.B and PHP.eB variants to cross the BBB and diffuse the brain parenchyma of mice ^36,37^. Interestingly, this receptor is not conserved in primates and humans, thus implying species-specific difference in the transduction efficacy of these capsids. On this line, new findings indicate that brain transduction in adult primates is not improved with PHP.B compared to AAV9 although only few animals have yet been tested^38,39^. Nonetheless, the PHP.B/eB receptor belongs to the large family of Ly6/uPar proteins, some of which are conserved in mammalian evolution and can be found in human brain endothelium, representing valuable targets for capsid engineering^40,41^. In this prospective, the PHP.B platform is fundamental to test the validity of new gene therapy strategies in mice models that can be translated into the clinical setting as soon as, in the next future, PHP.B-like AAVs penetrating the primate brain will be confirmed. Despite this caveat, RTT is a neurodevelopmental disorder and the therapeutic intervention should be finalized in the first months after birth as soon as the early signs of the disease manifest and genetic diagnosis is certain. At similar age, an intravenous infusion of AAV9 particles packaging the *SMN1* gene resulted in extended survival and improved motor functions in infants suffering for spinal muscular atrophy^42^. Thus, it is plausible that at this early age, AAV9 systemic gene therapy might sustain a beneficial clinical outcome in RTT patients and even more so using AAV9 synthetic variants selected in the near future. Altogether in this study we characterized an *iMecp2* viral cassette which sustained significant improvements in transgene delivery, safety and efficacy leading to the long-term symptomatic amelioration of RTT mice. These results might accelerate the introduction of new gene therapy strategies for RTT with clinical prospective.

## Materials and Methods

### Animals

Mice were maintained at San Raffaele Scientific Institute Institutional mouse facility (Milan, Italy) in micro-isolators under sterile conditions and supplied with autoclaved food and water. The *Mecp2*^-*/y*^ mice^4^ (The Jackson Laboratory stock *#003890*) were a kind gift of N. Landsberger and were maintained on C57BL/6J background. All procedures were performed according to protocols approved by the internal IACUC and reported to the Italian Ministry of Health according to the European Communities Council Directive 2010/63/EU.

### Generation of gene transfer vectors

All the Mecp2 and control AAV vectors were cloned into single stranded constructs. The murine *Mecp2* isoform 1 (NM_001081979.2) CDS including 3’-UTR (223bp) was PCR amplified in order to add the V5 tag at the 5’ of the coding sequence and inserted in the CBA-CreNLS vector to generate the M2a i*Mecp2* vector (Fig. 1a)^19^. The CBA promoter consists of 3 indipendent elements: a CMV enhancer, the chicken β-actin promoter and the human β-globin first intron. The CBA promoter was removed from the CBA-V5-*Mecp2* vector to be replaced by the mouse *Mecp2* promoter region (1,4Kb) including the 5’-UTR to generate the M2b vector (Fig. 1a). The AAV-CBA-V5-GFP construct was engineered from AAV-CBA-V5-Mecp2 vectors by exciding *Mecp2* CDS and exchanging with a GFP cassette. The 3’-UTR of the M2a construct was replaced by an assembled 3’-UTR (aUTR, 223bp, including portions of the 8,6Kb murine *Mecp2* endogenous 3’-UTR, such as: miRNA22-3p, 19-3p and 132-3p target sequence and the distal polyA signal of *Mecp2* gene)^15^ created by gene synthesis (Genewiz) or removed to generate respectively the M2c and M2e construct (Fig. 1a), whereas the M2-d construct was made from M2-a vector by insertion of the murine *Mecp2* 5’UTR upstream of the V5 sequence (Fig. 1a). In order to generate the iMECP2 vector the murine Mecp2 CDS and 3’-UTR of M2a was replaced by the human *MECP2* isoform_2 (NM_001110792.2) CDS including the human 3’-UTR (225bp) of *MECP2* gene. This isoform was chosen since is the human orthologue of murine *Mecp2* isoform_1. Both *NGFR* (Nerve Growth Factor Receptor) and *Mecp2* (comprehensive of 3’-UTR) amplicons were digested and cloned into a lentiviral vector (LV-Ef1a-GFP) in which the GFP cassette was removed.

### Virus production and purification

Lentiviral replication-incompetent, VSVg-coated lentiviral particles were packaged in 293T cells^43^. Cells were transfected with 30μg of vector and packaging constructs, according to a conventional CaCl_2_ transfection protocol. After 30 h, medium was collected, filtered through 0.44μm cellulose acetate and centrifuged at 20000 rpm for 2 h at 20°C in order to concentrate the virus.

AAV replication-incompetent, recombinant viral particles were produced 293T cells, cultured in Dulbecco Modified Eagle Medium – high glucose (Sigma-Aldrich) containing 10% fetal bovine serum (Sigma-Aldrich), 1% non-essential amino acids (Gibco), 1% sodium pyruvate (Sigma-Aldrich), 1% glutamine (Sigma-Aldrich) and 1% penicillin/streptomycin (Sigma-Aldrich). Cells were split every 3-4 days using Trypsin 0.25% (Sigma-Aldrich). Replication-incompetent, recombinant viral particles were produced in 293T cells by polyethylenimine (PEI) (Polyscience) co-transfection of three different plasmids: transgene-containing plasmid, packaging plasmid for rep and cap genes and pHelper (Agilent) for the three adenoviral helper genes. The cells and supernatant were harvested at 120 hrs. Cells were lysed in hypertonic buffer (40 mM Tris, 500 mM NaCl, 2 mM MgCl_2_, pH=8) containing 100U/ml Salt Active Nuclease (SAN, Arcticzymes) for 1h at 37°C, whereas the viral particles present in the supernatant were concentrated by precipitation with 8% PEG8000 (Polyethylene glycol 8000, Sigma-Aldrich) and then added to supernatant for an additional incubation of 30min at 37°C. In order to clarify the lysate cellular debris where separated by centrifugation (4000g, 30min). The viral phase was isolated by iodixanol step gradient (15%, 25%, 40%, 60% Optiprep, Sigma-Aldrich) in the 40% fraction and concentrated in PBS (Phosphate Buffer Saline) with 100K cut-off concentrator (Amicon Ultra15, MERCK-Millipore). Virus titers were determined using AAVpro^©^ Titration Kit Ver2 (TaKaRa).

### Primary mouse neuronal cultures

Primary Neuronal culture were prepared at embryonic day 17.5 (E17.5) from male mouse embryos. Briefly, cortices were individually dissected, sequentially incubated in trypsin (0,005%, 20 min at 37°C, Sigma-Aldrich) and DNAse (0,1mg/mL, 5min at room temperature, Sigma-Aldrich) in HBSS (Hank’s buffered salt solution without Ca^2+^ and Mg^2+^, Euroclone). Cells were finally and plated on poly-L-lysine (Sigma-Aldrich) coated dishes (2.0 × 10^5^ cells/cm^2^) in Neurobasal medium (TermoFisher Scientific) enriched with 0,6% glucose (Sigma-Aldrich), 0,2%penicillin/streptomycin (Sigma-Aldrich), 0,25% L-glutamine (Sigma-Aldrich) and 1% B27 (TermoFisher Scientific). Viral particles were directly added to cultured neurons 3 days after seeding, with a final concentration 10^10^ vg/ml.

### AAV-PHP.eB vector injection, mouse phenotyping and tissue collection

Vascular injection was performed in a restrainer that positioned the tail in a heated groove. The tail was swabbed with alcohol and then injected intravenously with a variable viral concentration (from 1*10^10^ to 1*10^13^ vg/mL) depending on the experimental setup in a total volume of 100 µl of AAV-PHP.eB particles in PBS.

Juvenile WT *and Mecp2*^−/y^ mice were randomized in groups and injected in the tail vein between 25 and 30 days of age. Adult WT were injected in a similar time window whereas *Mecp2^+/-^* females were treated intravenously after five months of life. Following injection, all mice were weighed twice a week. Phenotyping was carried out, blind to genotype and treatment, twice a week. Mice symptoms were scored on an aggregate severity scale (0 = absent; 1 = present; 2 = severe) comprising mobility, gait, breathing, hindlimb clasping, tremor, and general condition. The balance and the motor coordination were assessed by the Beam Balance test. Briefly mice were placed on the tip of the beam at the “start-point” facing towards the beam. The number of foot-slips (error numbers) and total time on beam (crossing time) from “start” to “end” points were noted. If a mouse fell, the animal was returned to the site where it fell from, until completion of beam crossing. The Open Field test was performed in an arena of 50 × 50 cm. Mice were testing in a 10 minutes session measured as an index of anxiety and horizontal exploratory activity in a novel environment was assessed.

For serum analysis blood samples were collected from living animals using retro-orbital bleeding procedure with non-heparinized capillaries. Upon blood clothing cell fraction was pelleted (5min, 13000 rpm) and supernatant recovered. When the body loss reached 20% of total weight mice were sacrificed and tissues harvested. Briefly, mice were anesthetized with ketamine/xylazine and transcardially perfused with 0.1 M phosphate buffer (PB) at room temperature (RT) at pH 7.4. Upon this treatment brain, liver and spleen were collected. Brain hemispheres were separated: one half was post-fixed in 4% PFA for two days and then soaked in cryoprotective solution (30% sucrose in PBS) for immunofluorescence analysis the other further sectioned in different areas (cortex, striatum. cerebellum) quick frozen on dry-ice for Western blot, RNA and DNA extraction. Liver specimens were collected similarly. Spleens were collected in PBS for subsequent splenocyte extraction.

### Total RNA-DNA isolation and qRT-PCR for Mecp2 RNA stability, biodistribution and gene expression

Total RNA was isolated from primary neurons and animal tissues (cortex and liver) using the Qiagen RNeasy mini kit (QIAGEN). About 1µg of RNA was reverse transcribed with random hexamers as primers using ImProm-II™ Reverse Transcription System (Promega). For quantitative real time PCR (qRT-PCR), Titan HotTaq EvaGreen qPCR mix (BioAtlas) was used and expression levels were normalized respect to β-actin expression. The results were reported as the fold change (2^-ΔΔCt^) of viral *Mecp2* relative to endogenous *Mecp2*.

The stability of endogenous and viral *Mecp2* RNA was assessed by qRT-PCR (**Supplementary Table 2**). The RNA was isolated at the indicated time-points from neurons (WT uninfected and infected with *iMecp2* vector and *Mecp2^-/y^* infected with *iMecp2* vector) treated with 10 mg/mL of Actinomycin D (Sigma-Aldrich).

Total DNA was isolated from primary neurons and animal tissues (cortex and liver) using the Qiagen DNeasy Blood & Tissue Kits (QIAGEN). The quantification of vector transgene expression was calculated by qRT-PCR relative to the endogenous *Mecp2*. The DNA levels were normalized against an amplicon from a single-copy mouse gene, *Lmnb2*, amplified from genomic DNA.

### RiboLace

The RiboLace kit (IMMAGINA Biotechnology S.r.l) was used to isolate from primary neurons (*wild-type* uninfected and infected with *iMecp2* vector and *Mecp2^-/y^* infected with *iMecp2* vector) at DIV 14 two distinct fraction: total RNA and RNA associated to the active ribosomes, according to manufacturer’s instructions. Following the isolation, about 100 ng of RNA was reverse transcribed and amplified by qRT-PCR as described above using the oligonucleotide primers to amplify endogenous *Mecp2*, viral *Mecp2* and 18S as housekeeping gene (**Supplementary Table 2**). The result was reported as fold change (2^-ΔΔCt^) in gene expression of viral *Mecp2* relative to endogenous *Mecp2*, in the captured fraction normalized on total RNA.

### CHX-chase analysis

Primary neurons (WT uninfected and *Mecp2^-/y^* infected with the *iMecp2* vector) were incubated with CHX (50µM). The cells were collected after 0 and 8 hours after treatment for protein extraction. Lysate samples were finally prepared for Western blot analysis as described below.

### Generation of a MECP2-KO human iPS cell (iPSC) line

Control human iPSC cell line were generated from neonatal primary fibroblasts obtained from ATCC. iPSCs were maintained in feeder-free conditions in mTeSR1 (Stem Cell Technologies) and seeded in HESC qualified matrigel (Corning)-coated 6-well plates. To generate the MECP2-KO cell line, an sgRNA (sgMECP2: 5’-aagcttaagcaaaggaaatc-3’) was designed on the third exon of MECP2 using the software *crispor.tefor.net*. The oligo (Sigma-Aldrich) pairs encoding the 20-nt guide sequences were cloned into the LV-U6-filler-gRNA-EF1α-Blast (Rubio et al., 2016). Wild-type human iPSCs were then co-transfected with the LV-U6-sgMECP2-EF1α-Blast and the pCAG-Cas9-Puro using the Lipofectamine Stem Cells Transfection Reagent (ThermoFisher Scientific)^44^. Co-transfected colonies were then selected by the combination of puromycin (1µg/ml, Sigma) and blastidicin (10 μg/ml, ThermoFisher Scientific) and then isolated through single colony picking. Finally, MECP2-KO cell lines were confirmed by Sanger Sequencing and protein absence was further corroborated by immunofluorescence.

### Differentiation of human iPSCs in cortical neurons

iPSCs were initially differentiated in Neural Progenitors Cells (NPCs) as described in Iannielli et al^45^NPCs were, then, dissociated with Accutase (Sigma-Aldrich) and plated on matrigel-coated 6-well plates (3*10^5^ cells per well) in NPC medium. Two days after, the medium was changed with the differentiation medium containing Neurobasal (ThermoFisher Scientific), 1% Pen/Strep (Sigma-Aldrich), 1% Glutamine (Sigma-Aldrich), 1:50 B27 minus vitamin A (ThermoFisher Scientific), 5 μM XAV939 (Sigma-Aldrich), 10 μM SU5402 (Sigma-Aldrich), 8 μM PD0325901 (Tocris Bioscience), and 10 μM DAPT (Sigma-Aldrich) was added and kept for 3 days. After 3 days, the cells were dissociated with Accutase (Sigma-Aldrich) and plated on poly-L-lysine (Sigma-Aldrich)/laminin (Sigma-Aldrich)-coated 12-well plates (2*10^5^ cells per well) and 24-well plates (1*10^5^ cells per well) in maturation medium containing Neurobasal (ThermoFisher Scientific), 1% Pen/Strep (Sigma-Aldrich), 1% Glutamine (Sigma-Aldrich), 1:50 B27 minus vitamin A (ThermoFisher Scientific), 25 ng/ml human BDNF (PeproTech), 20 μM Ascorbic Acid (Sigma-Aldrich), 250 μM Dibutyryl cAMP (Sigma-Aldrich), 10 μM DAPT (Sigma-Aldrich) and Laminin for terminal differentiation. At this stage half of the medium was changed every 2–3 days. Viral particles were directly added to cultured neurons after three weeks of differentiation, with a final concentration 10^10^ vg/ml. All the analysis was conducted one week after the infection.

### Immunofluorescence

Primary neurons and human iPSCs-derived neurons were fixed with ice-cold 4% paraformaldehyde (PFA) for 30 min at 4°C, washed with PBS (3×) and incubated with 10% donkey serum and 3% Triton X-100 for 1 hr at RT to saturate the unspecific binding site before the overnight incubation at 4°C with the primary antibody. Upon wash with PBS (3×), cells were incubated for 1 h at RT in blocking solution with DAPI and with Alexa Fluor-488 and Alexa Fluor-594 anti-rabbit or anti-mouse secondary antibodies. After PBS washes (3×), cells were mounted with fluorescent mounting medium (Dako). Images were captured with a Nikon Eclipse 600 fluorescent microscope.

Tissues were sectioned using cryostat after optimal cutting temperature compound (OCT) embedding in dry ice. Free-floating 50μm-thick coronal sections were rinsed in PBS and were incubated with 10% donkey serum (Sigma-Aldrich) and 3% Triton X-100 (Sigma-Aldrich) for 1 hr at RT to saturate the unspecific binding site before the overnight incubation at 4°C with the primary antibody (diluted in the blocking solution).

Upon wash with PBS (3×), sections were incubated for 1 h at RT in blocking solution with DAPI (1:1000, Sigma-Aldrich) and with Alexa Fluor-488 and Alexa Fluor-594 anti-rabbit or anti-mouse secondary antibodies (1:1000, ThermoFisher Scientific). After PBS washes (3×), sections were mounted with fluorescent mounting medium (Dako). Confocal images were captured at ×40 or ×63 magnification with Leica TCS SP5 Laser Scanning Confocal microscope (Leica Microsystems Ltd). Cell and tissue where stained with the following primary antibody: rabbit anti-MeCP2 (1:500; Cell Signaling Technology), mouse anti-V5 (1:500; ThermoFisher Scientific), mouse anti-NeuN (1:300; Merck Millipore), rabbit anti-GFAP (1:500; Dako), rabbit anti-GABA (1:500; Sigma-Aldrich) anti-chicken_MAP2 (1:500, Abcam), anti-OCT4 (1:50, Abcam).

### Western blot

Protein extracts were prepared in RIPA buffer (10 mM Tris-HCl pH7.4, 150 mM NaCl, 1 mM EGTA, 0.5% Triton and complete 1% protease and phosphatase inhibitor mixture, Roche Diagnostics). Primary neurons, brain and liver lysate samples (50 μg protein lysates) were separated using 8% polyacrylamide gel and then transferred to PVDF membranes. Membranes were incubated overnight at 4°C with the following primary antibodies in 1X PBST with 5% w/v nonfat dry: rabbit anti-MeCP2 (1:1000; Sigma), mouse anti-V5 (1:1000; ThermoFisher Scientific), rabbit anti-pS6 235/236 (1:500, Cell Signaling), rabbit anti-S6 (1:500, Cell Signaling), rabbit anti-Calnexin (1:50000, Sigma), mouse anti-β-Actin (1:50000; Sigma) or the mice serum (1:200) extracted through retro-orbital bleeding followed by centrifugation (10 min, RT, 13000 rpm). Subsequently, membranes were incubated with the corresponding horseradish peroxidase (HRP)-conjugated secondary antibodies (1:10000; Dako). The signal was then revealed with a chemiluminescence solution (ECL reagent, RPN2232; GE Healthcare) and detected with the ChemiDoc imaging system (Bio-Rad).

### Antibody detection in serum

Serum was extract from mice through retro-orbital bleeding followed by centrifugation (10 min, RT, 13000 rpm). In order to test the sera by immunofluorescence we generated a P19 *Mecp2^-/y^* cell line using pCAG-spCas9^44^ and sg*Mecp2*^46^. Moreover, isolation of a clone carrying a frameshift mutation (−14nt) in the exon 3 of *Mecp2* gene that ensured ablation of MeCP2 protein (as tested by immunofluorescence). *Mecp2^-/y^* P19 cells were transfected with GFP only (negative control) or with GFP and *iMecp2*, cells were fixed and incubated in blocking solution as describe above before the overnight incubation at 4°C with the primary antibody mix composed of chicken anti-GFP and either a rabbit anti-MeCP2 (positive control) or the mice serum (1:50) (for details see the above Immunofluorescence paragraph). For Western blot assay, the proteins *Mecp2^-/y^* and WT cortices were extracted and separated using 8% polyacrylamide gels. Membranes were incubated overnight at 4°C with a rabbit anti-MeCP2 (positive control) or the mice serum (1:200) (for details see the Western blot paragraph).

### Fluorescent intensity measurements

Brain sections were processed for immunolabeling as above and confocal images were captured at ×63 magnification with Leica TCS SP5 Laser Scanning Confocal microscope (Leica Microsystems Ltd) using the identical settings. Then, the quantification of the signal was performed using ImageJ software (NIH, US). The fluorescent signal was measured as described^45^.

### Spleen cell isolation

Spleens were triturated in PBS and cell pelleted (7min, 1500rpm) to be incubated in ACK buffer (5min, RT) to lyse blood cells. The reaction was stopped diluting 1:10 the ACK buffer in PBS and removing it by centrifugation (7min, 1500rpm). Cells were than counted and frozen in FBS:DMSO (9:1 ratio) solution.

### Flow cytometry

Spleen cells were incubated with 25 µl of Ab mix listed in Supplementary Table 3 for 30’ at 4°. Red blood cells lysis was performed with BD Phosflow (BD Bioscience, 558049) according to manufacturer’s instruction. Labelled cells were washed two times with PBS 1% FBS and analysed with a BD LSRFortessa analyser, results were analysed with FlowJo 10 software.

### T cell proliferation

Spleen cells were labelled with Cell Proliferation Dye eFluor® 670 (eBioscience, CA, USA) according to manufacturer’s instructions and stimulated with 10^4^ bone-marrow derived DC transduce with lentiviral vector encoding for *Mecp2* (10:1, T:DC) in RPMI 1640 medium (Lonza, Switzerland), with 10% FBS (Euroclone, ECS0180L), 100 U/ml penicillin/streptomycin (Lonza, 17-602E), 2 mM L-glutamine (Lonza, 17-605E), Minimum Essential Medium Non-Essential Amino Acids (MEM NEAA) (GIBCO, 11140-035), 1 mM Sodium Pyruvate (GIBCO, 11360-039), 50 nM 2-Mercaptoethanol (GIBCO, 31350-010). Alternatively, spleen cells were stimulated with anti-CD3e monoclonal Ab (BD Bioscience, 553058) (1 μg/mL). After 4 days, T cells were collected, washed, and their phenotype and proliferation were analysed by flow cytometry. Antibodies used for flow cytometry are listed in Supplementary Table 3.

### Elispot Assays

CD8^+^ T cells were magnetically isolated from the spleen (Miltenyi Biotec, 130-104-075). 10^5^ CD8^+^ T cells were plated in triplicate in ELISPOT plates (Millipore, Bedford, MA) pre-coated with anti– IFN-γ capture monoclonal Ab (2.5 μg/mL; BD Pharmingen, R46A2) in the presence of IL-2 (50 U/mL; BD Pharmingen) and 10^5^ irradiated (6000 rad) un-transduced or LV.*Mecp2*-transduced autologous EL-4 cells. After 42 hours of incubation at 37°C 5% CO_2_, plates were washed and IFN-γ–producing cells were detected by biotin-conjugated anti–IFN-γ monoclonal Ab (0.5 μg/mL; BD Pharmingen, XMG 1.2). Streptavidin-HRP conjugate (Roche) was added. Total splenocytes or total BM (0,35 x 10^6^ cells/well) were plated in complete RPMI in triplicate in ELISPOT plates pre-coated with rhIDUA (2µg/well). After 24 hours of incubation at 37°C 5% CO_2_, plates were washed and anti-IDUA IgG secreting cells were detected with peroxidase-conjugated rabbit anti–mouse immunoglobulin (SIGMA A2554). All plates were reacted with H_2_O_2_ and 3-Amino-9-ethylcarbazole (SIGMA, A6926). Spots were counted by ImmunoSpot reader (Cellular Technology Limited)

### Computational analysis

FASTQ reads were quality checked and trimmed with FastQC^47^. High quality trimmed reads were mapped to the mm10/GRCm38.p6 reference genome with Bowtie2 v2.3.4.3^48^. Gene counts and differential gene expression were calculated with featureCount using latest GENCODE main annotation file^49^ and DESeq2^50^ respectively. Geneset functional enrichment was performed with GSEA^51^. Downstream statistics and Plot drawing were performed with R. Heatmaps were generated with GENE-E (The Broad Institute of MIT and Harvard).

### Rotarod

Mice were assessed on an accelerating rotarod (Ugo-Basile, Stoelting Co.). Revolutions per minute (rpm) were set at an initial value of 4 with a progressive increase to a maximum of 40 rpm across the 5 minutes test session. The animals were given two days of training and one day of test, each session consisting of three trials. Latency to fall was measured by the rotarod timer.

### Elevated Plus Maze (EPM)

The test uses an elevated cross (+) apparatus with two open and two enclosed arms (length 45 cm): the open arms had low walls (0,5 cm) while the closed arms had high walls (20 cm). The mouse was placed in the center of the apparatus and it could move and explore freely the maze for 10 minutes. The amount of the time that the animal spent in closed arms were measured and analyzed with EthoVision XT system. The maze was cleaned with water and 70% ethanol before the next mouse was placed on the apparatus.

### Statistics

Values are expressed as mean ± standard deviation as indicated. All statistical analysis was carried out in GraphPad Prism 8.0, using one-way ANOVA, two-way ANOVA, Mantel-Cox test (survival curves) and non-parametric Mann-Whitney U test (two-tailed) where unpaired t-test was applied. P-values below 0.05 were considered significant. In multi-group comparisons, multiple testing correction for pairwise tests among groups was applied using Tukey’s post hoc analysis.

## Acknowledgements

We thank N. Landsberger, E. Cattaneo, F. Ciceri and L. Naldini for providing valuable reagents and mouse strains. We are grateful to D. Bonanomi, S. Biffo and all members of the Broccoli’s lab for helpful discussion. We acknowledge the FRACTAL and ALEMBIC core facilities for expert supervision in flow-cytometry and confocal imaging, respectively. This work was supported by the Telethon Foundation grant (GGP19038) to V.B.

## Author contributions

M.L. and S.Gi. performed the experiments and analyzed data; M.I. carried out and analyzed most of the behavioral studies; A.N. performed molecular cloning and immunoblotting; L.M. developed the computational analysis; L.P and S.Gr. performed and analyzed immune response in animal tissues; P.C. contributed to perform and analyze RT-qPCR assays; G.M. executed tail vein injections and brain histopathological analysis; B.D contributed in designing the experiments; V.B. supervised, coordinated and supported the project and wrote the paper with M.L., S.Gi. and S.Gr.

## Supplementary Materials

**Supplementary Figure 1. Distribution and quantification of *iMecp2* gene transfer in *Mecp2^-/y^* brains.**

(**a**) Low magnification of Mecp2 immunostaining of forebrain (upper panel) and cerebellum (lower panel) sections in KO control untreated (UT) and treated animals (1*10^10^, 1*10^11^, 1*10^12^ vg/mouse). (**b**) High magnification confocal images for Mecp2 and V5 in cortex derived from WT and *iMecp2* treated KO (untreated [WT], 1*10^11^ and 1*10^12^ vg/mouse) animals. Nuclei were stained with DAPI. Scale bar: 10μm. Left panel: graphs showing the quantification of cellular levels of total Mecp2 detected with anti-MeCP2 immunofluorescence in cells of the cortex and quantified in arbitrary units based on field pixel intensity (n=30 nuclei for per condition, UN: untransduced). Error bars, SD. *** p < 0.001 as compared to untreated (WT) (ANOVA-one way, Tukey’s post hoc test). Scale bars: 500 µm (a), 10 µm (b).

**Supplementary Figure 2. Cell type analysis of *iMecp2* gene transfer in *Mecp2^-/y^* brains.**

(**a**) Characterization of *iMecp2*-transduced cells in KO cortex (1*10^11^ vg/mouse) using colocalization of V5^+^ cells with markers of brain cellular sub-populations such as: neurons (marked with NeuN, upper panel), astrocytes (marked with GFAP, middle panel), and GABAergic interneurons (marked with GABA, lower panel). Nuclei were stained with DAPI. All images were captured using confocal microscope. Scale bar: 50μm. (**b**) Bar graphs showing co-localization of these markers with V5 or Mecp2 (n = 3) (**c**) High magnification confocal images of wild-type (WT, upper panel) and *iMecp2* transduced nuclei (lower panel) stained with MeCP2 antibody both exhibit heterochromatin-enriched localization. Error bars, SD. Scale bars: 50 µm (a), 5 µm (c).

**Supplementary Figure 3. Characterization of *iMecp2* transgene expression in brain and liver of *Mecp2^-/y^* animals.**

(**a**) Vector biodistribution (upper panel) and transgene expression (lower panel) in mice cortex and liver of KO mice untreated (KO, n = 3) and treated with 1*10^11^ (n = 3) or 1*10^12^ (n = 3) *iMecp2*. Data were normalized over *Mecp2* gene levels and expression of wild-type mice (WT, light blue bar), respectively. (**b**) Western blot analysis to quantify Mecp2 over Actin protein levels in liver (left panel) from WT, untreated and *iMecp2* treated (1*10^10^, 1*10^11^, 1*10^12^ vg/mouse) KO animals and corresponding densitometric quantification expressed in arbitrary units (n = 5 for 1*10^10^, n = 7 for 1*10^11^; n =5 for 1*10^12^ vg/mouse). Error bars, SD. *** p < 0.001 as compared wild-type (WT) (ANOVA-one way, Tukey’s post hoc test).

**Supplementary Figure 4. Behavioral rescue of symptomatic *Mecp2^-/y^* animals after PHP.eB-*iMecp2* treatment.**

(**a**) General phenotypic assessment evaluated through the aggregate severity score (p < 0.05 versus KO-GFP in 1*10^10^ [7^th^-10^th^ wk] and in 1*10^11^ [6^th^-10^th^ wk]). (**b**) Mouse body weight was monitored every two weeks and represented as mean of each group (p < 0.05 versus KO-GFP in 1*10^10^ [5^th^-10^th^ week, wk] and in 1*10^11^ [7^th^-9^th^ wk]). (**c**) Spontaneous locomotor activity was tested in a spontaneous field arena and shown as representative traces and quantification of total distance. (** p < 0.01 and *** p < 0.001 as compared to WT mice). (**d**) Representative pictures of animals WT, KO and KO treated with most efficacious dose (1*10^11^) indicate the absence of hindlimb clasping in KO treated animals up to 14 weeks of age. (**e,f**) All groups of animals were tested for motor coordination using beam balance test and quantified as crossing time (**e**, p < 0.05 versus KO-GFP in 1*10^10^ [7^th^-10^th^ wk] and in 1*10^11^ [7^th^-9^th^ wk]) and number of errors (**f,** p < 0.05 versus KO-GFP in 1*10^10^ [8^th^-10^th^ wk] and in 1*10^11^ [7^th^-10^th^ wk]). Error bars, SD. ANOVA-two way (b, e, f, g) or ANOVA-one way (c) with Tukey’s post hoc test.

**Supplementary Figure 5. PHP.eB-*iMecp2* gene transfer in *Mecp2^+/-^* females.**

(**a**) Illustration of the experimental setting to restore the expression of Mecp2 in heterozygous (Het) animals by means of AAV-PHP.eB. (**b**) Mouse body weight was monitored every two weeks and represented as mean of each group. (**c**) General phenotypic assessment evaluated through aggregate severity score. (**d**) Spontaneous locomotor activity was tested in a spontaneous field arena and shown as quantification of travelled total distance. All groups of animals were tested for motor coordination using beam balance test and quantified as crossing time (**e**) and number of errors (**f**) (WT untreated [n=8], Het treated with GFP 1*10^11^ vg/mouse [n = 6], KO treated with *iMecp2* virus 1*10^11^ vg/mouse [n = 6]). Error bars, SD. ** p < 0.01. ANOVA-two way (b, c, e, f) or ANOVA-one way (d) with Tukey’s post hoc test.

**Supplementary Figure 6. PHP.eB-mediated *iMecp2* gene transfer in symptomatic *Mecp2^+/-^* mouse brains.**

(**a**) High magnification immunostaining for Mecp2 and V5 in cortex, striatum and cerebellum derived from WT, *Mecp2^+/-^* (Het) untreated and Het treated with PHP.eB*-iMecp2* (1*10^11^ vg/mouse) brains. Nuclei were stained with DAPI (merge panels). Bar graphs showing the fraction of V5 positive on the total DAPI positive in cortex, striatum and cerebellum (n = 3). (**b**) Western-blot analysis to quantify Mecp2 over Actin protein levels in cortex derived from WT, Het untreated and treated with *iMecp2* (1*10^11^) animals and corresponding densitometric quantification expressed in arbitrary units (right panel) (n = 3 for WT, n = 3 Het untreated; n = 3 Het treated with PHP.eB*-iMecp2* 10^11^ vg/mouse) * p < 0.001 as compare to WT and Het treated with PHP.eB*-iMecp2* (ANOVA-one way, Tukey’s post hoc test).

**Supplementary Figure 7. Characterization of *iMecp2* transgene expression in brain and liver of wild-type animals.**

Vector biodistribution (upper panel) and transgene expression (lower panel) in mice cortex and liver of WT mice untreated (UT, n = 3) and treated with 1*10^11^ (n = 3) or 1*10^12^ (n = 3) *iMecp2*. Data were normalized respectively over *Mecp2* gene level and expression of WT mice (UT, light blue bar). Error bars, SD.

**Supplementary Figure 8. Protein levels in PHP.eB-iMecp2 transduced wild-type animals.**

(**a**) Western blot analysis to quantify Mecp2 over Actin protein levels in striatum (upper panel) and liver (lower panel) derived from WT untreated (UT) and treated with *iMecp2* (1*10^11^ and 1*10^12^ vg/mouse) mice. Corresponding densitometric quantification expressed in arbitrary units (n = 3 for UT, n = 3 for 1*10^11^; n = 3 for 1*10^12^ vg/mouse) are shown on the right. (**b**) High magnification confocal images for V5 and Mecp2 in cortex derived from wild-type untreated (WT) and treated with *iMecp2* (1*10^11^, 1*10^12^ vg/mouse) animals. Nuclei were stained with DAPI. Scale bar: 10μm (Right panel). Left panel: graphs showing the quantification of cellular levels of total Mecp2 detected by immunofluorescence in cells of the cortex and quantified in arbitrary units based on field pixel intensity (n = 30 nuclei for per condition, UT: untreated). Error bars, SD. ** p < 0.05 and *** p < 0.001 compared to UT (ANOVA-one way, Tukey’s post hoc test). Scale bar: 10 µm.

**Supplementary movie. Activity of *Mecp2^−/y^* treated with GFP or 1*10^11^ vg/mouse *iMecp2* + cyclosporine.**

Both KO mice were injected after 4 weeks from birth and were video-recorded after 3 weeks (GFP, 1*10^11^ vg/mouse) or 4 months (*iMecp2*, 1*10^11^ vg/mouse) from the treatment. The *iMecp2*-treated mouse walks, climbs, explores and interacts with the other mouse, while the mock GFP-treated KO mouse remains stationary at one corner of the cage. This video indicates an improved mobility and an increased sociability after gene therapy in KO mice

**Supplementary Table 1: No evidence of liver toxicity in mice administered with high AAV dose.**

Blood serum levels of liver enzyme and liver histochemical analysis (representative images) were used as indicators of liver health. # Reference values for C57BL/6J male mice were taken from the mouse phenome database https://phenome.jax.org/. Abbreviations, Treat: treatment, ALB: albumin, ALP: Alkaline phosphatase, ALT: Alanine aminotransferase, HE: hematoxylin/eosin staining. All values are indicated as mean ± SD, n = 3, SD. Scale bar: 200 µm

**Supplementary Table 2: Primers employed for qRT-PCRs.**

**Supplementary Table 3: List of antibodies for flow cytometry.**

