## Supplementary data for "Whole brain delivery of an instability-prone *Mecp2* transgene improves behavioral and molecular pathological defects in mouse models of Rett syndrome"

#### Supplementary Figure 1

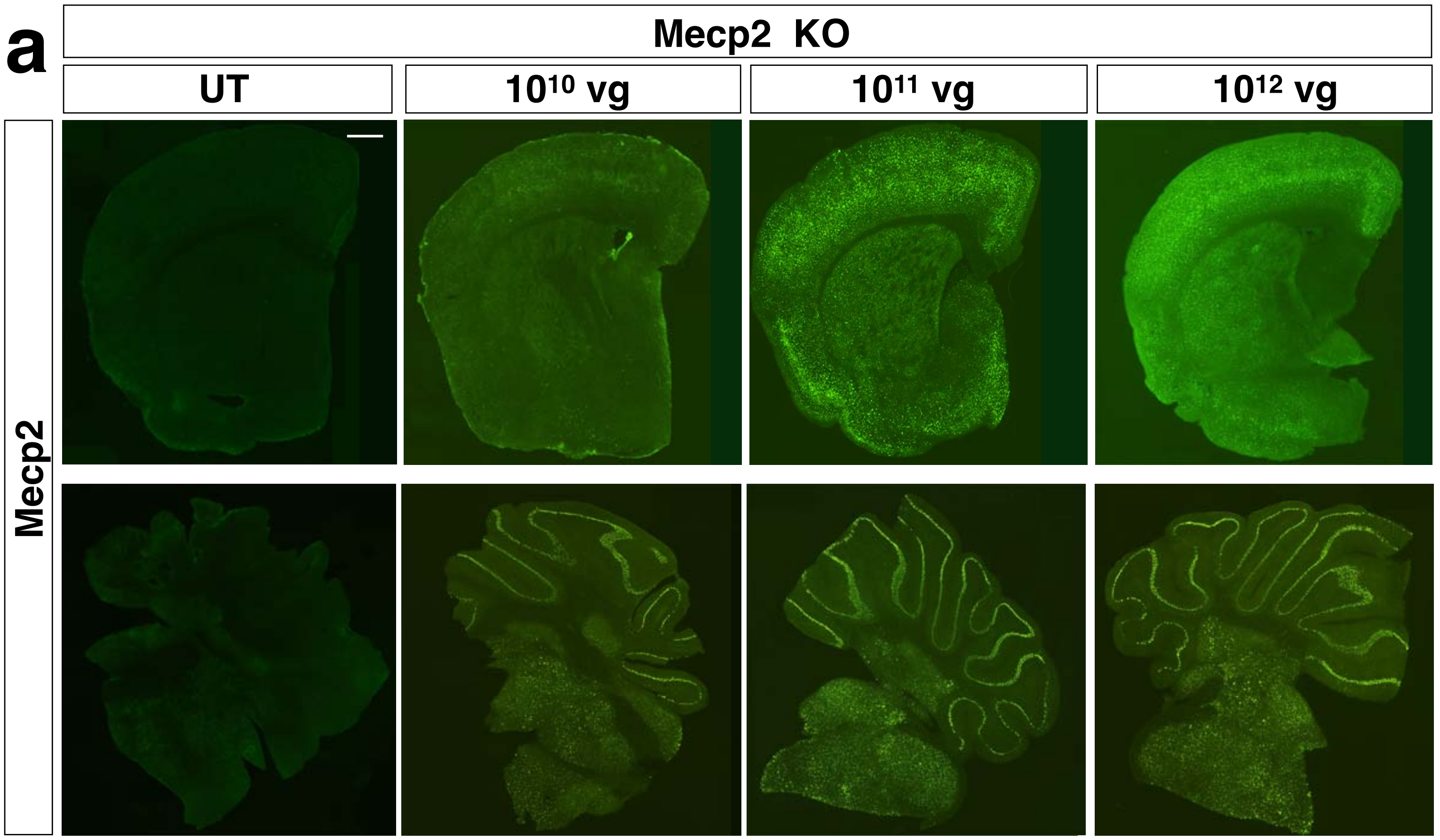


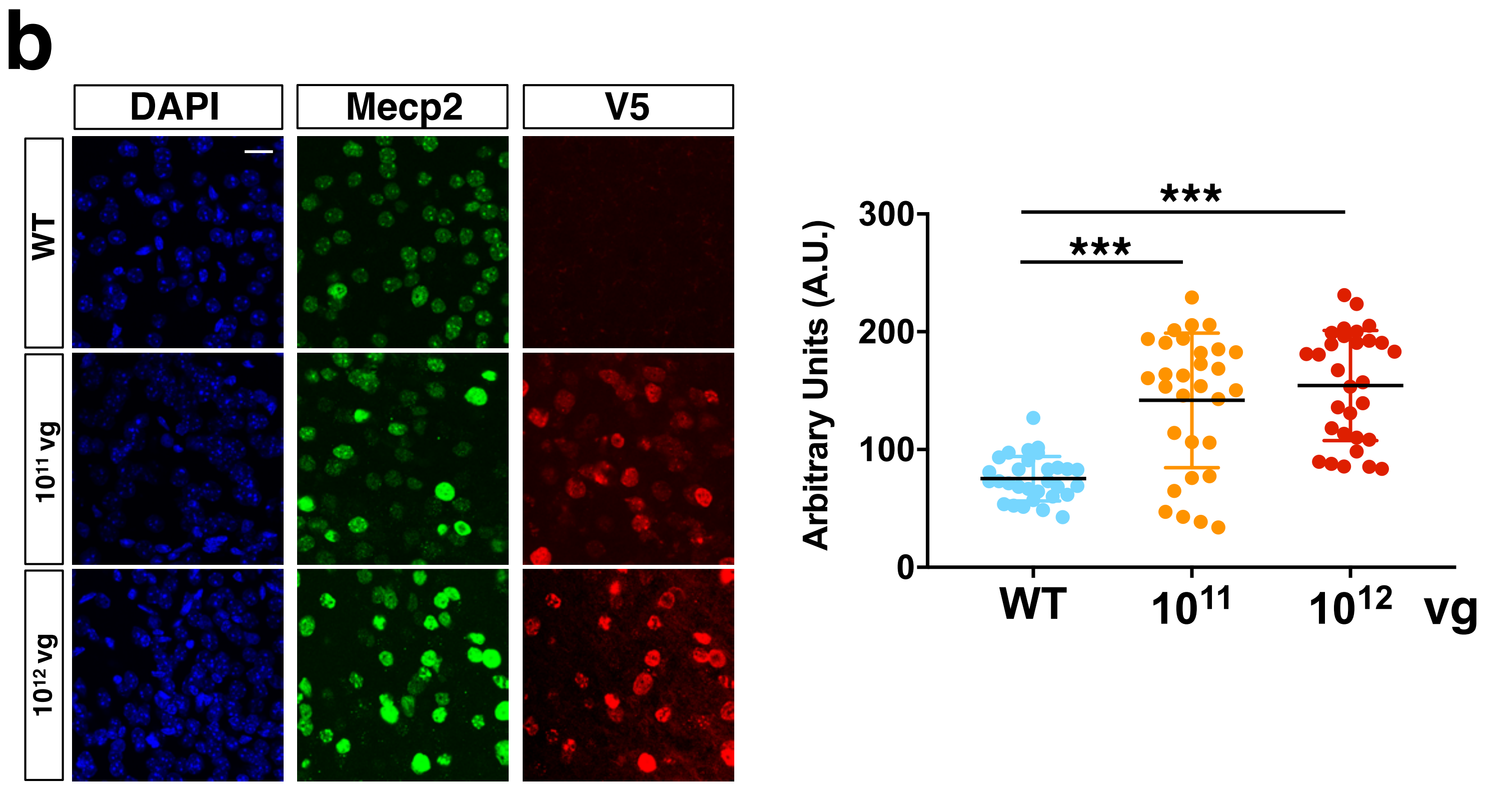

### Supplementary Figure 2

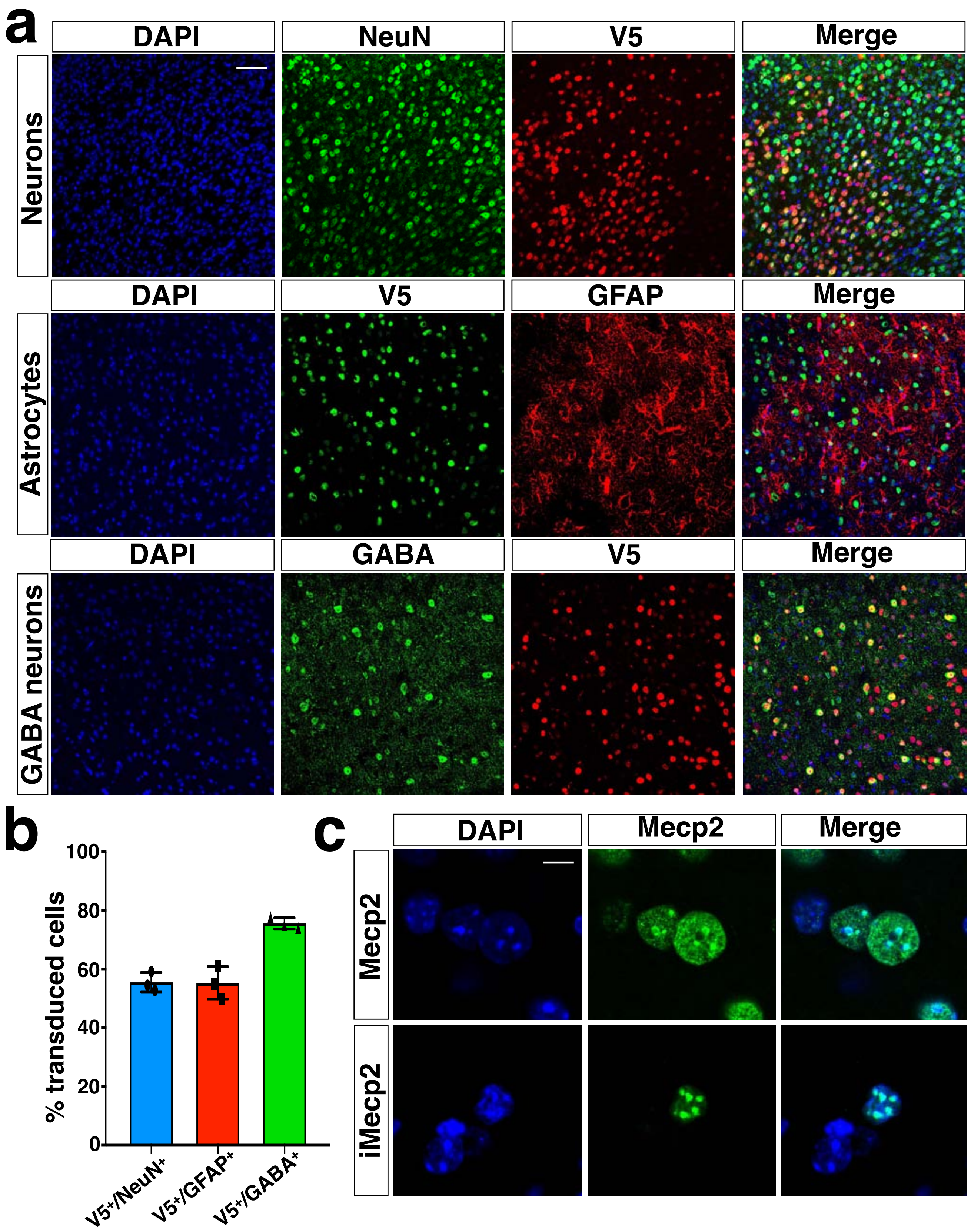

#### Supplementary Figure 3

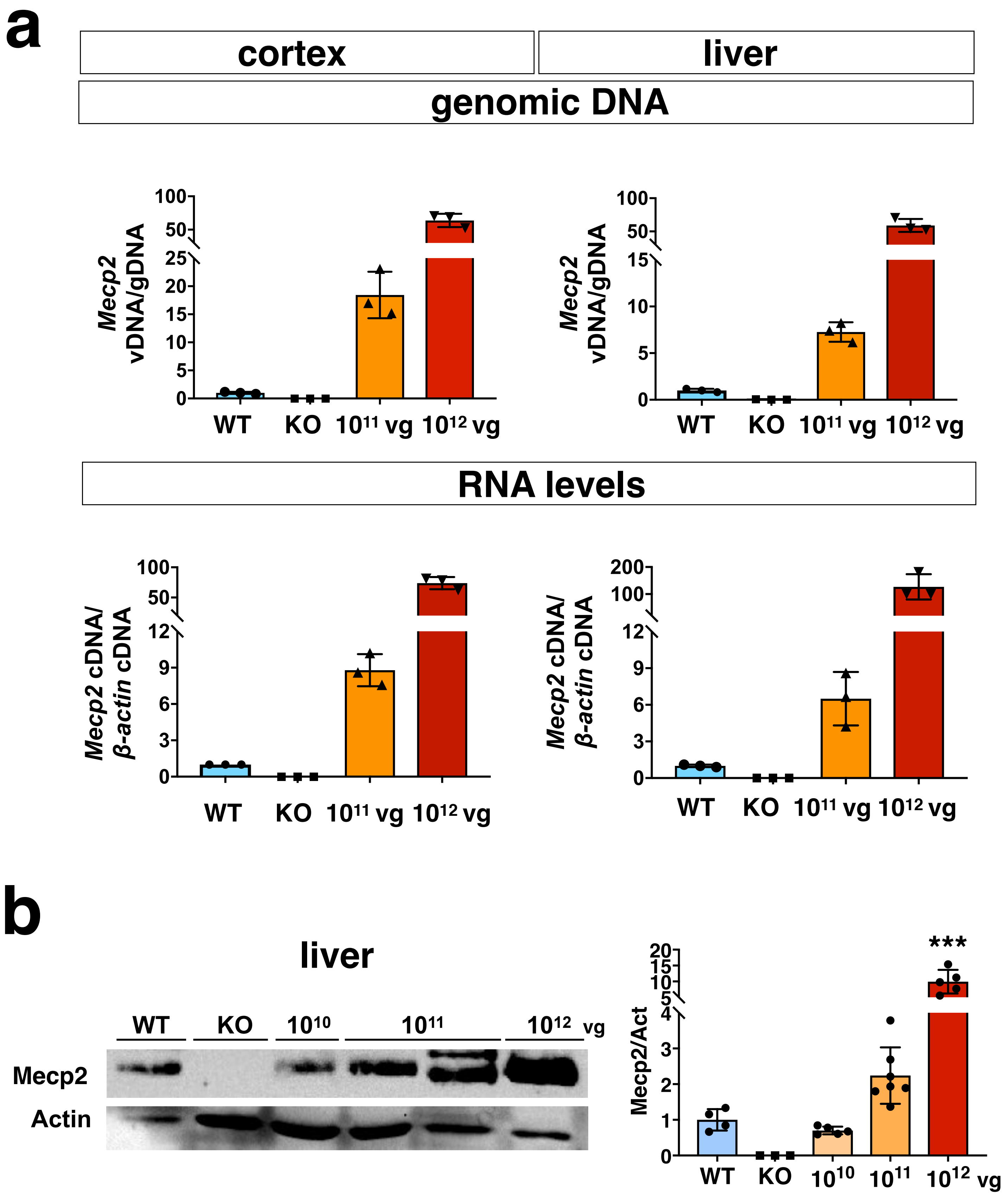

#### Supplementary Figure 4

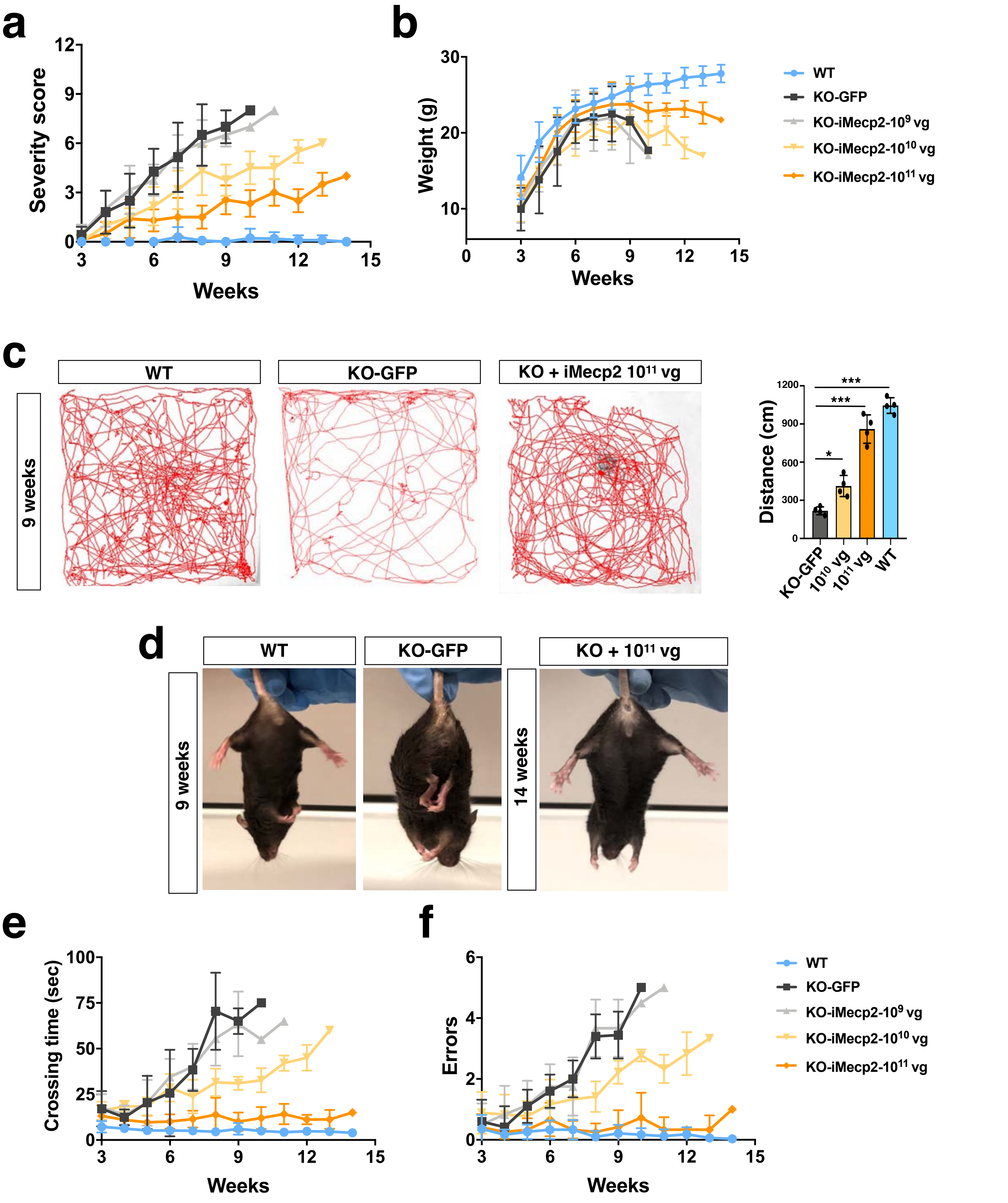

#### Supplementary Figure 5

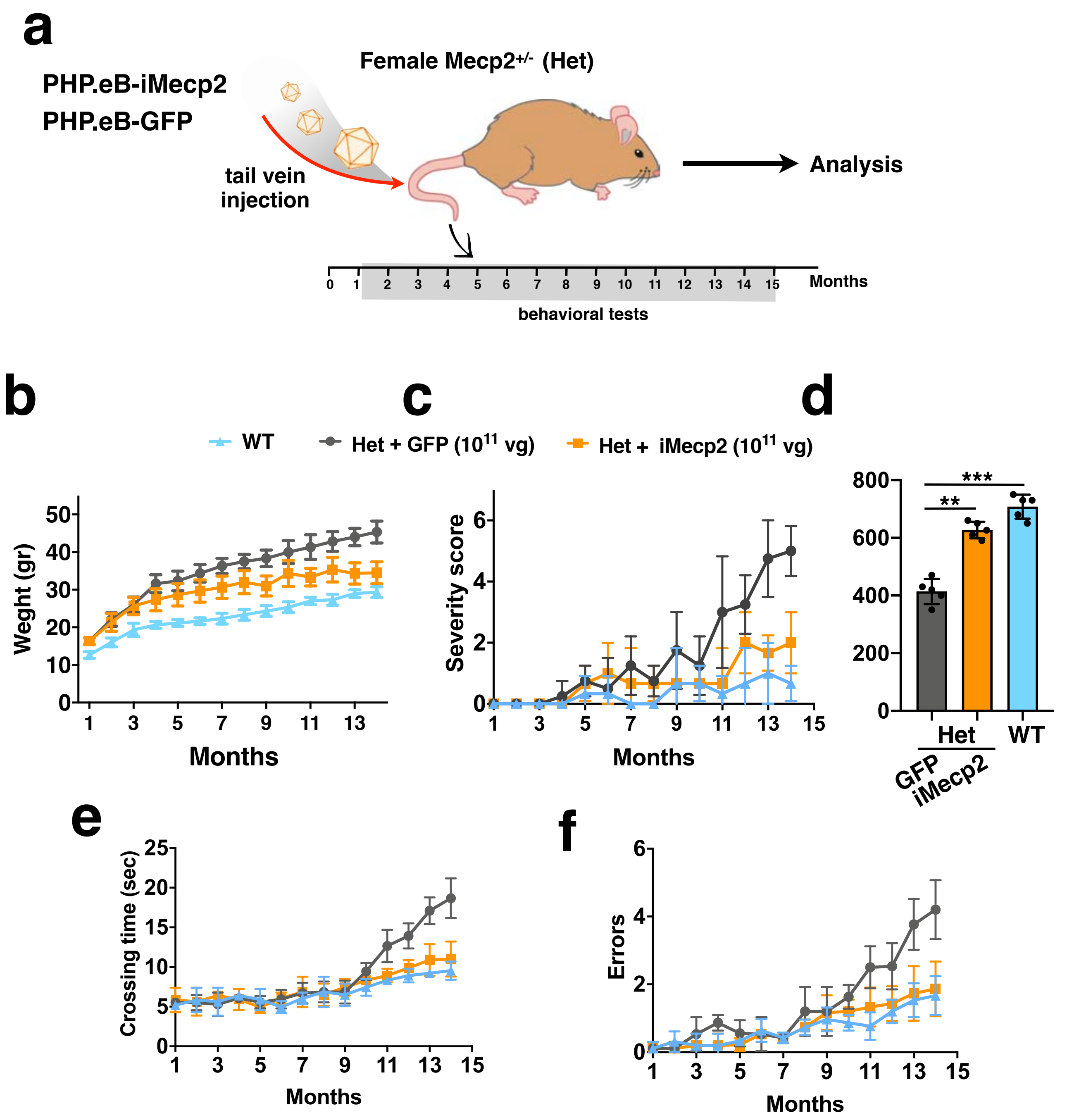

#### Supplementary Figure 6

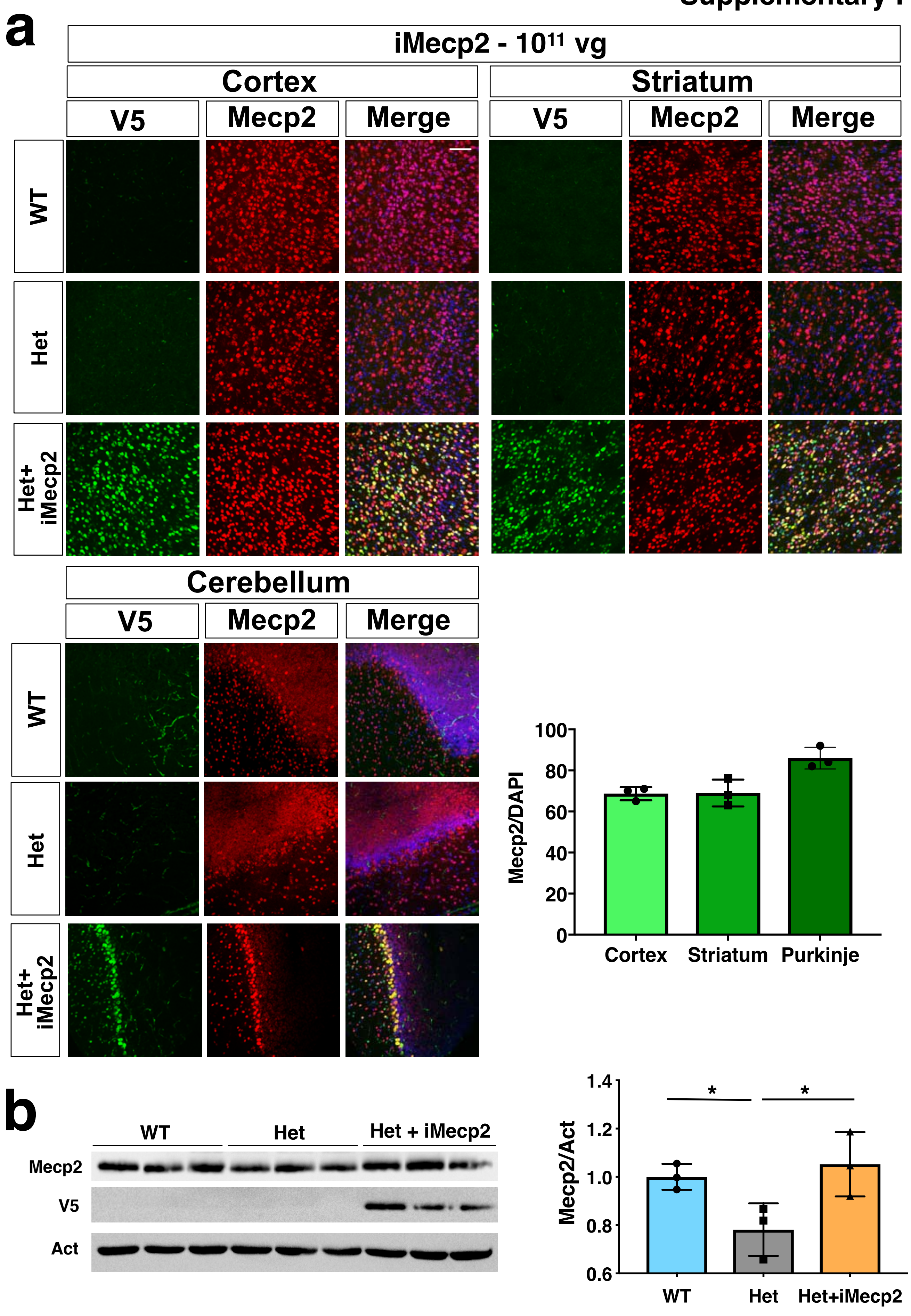

#### Supplementary Figure 7

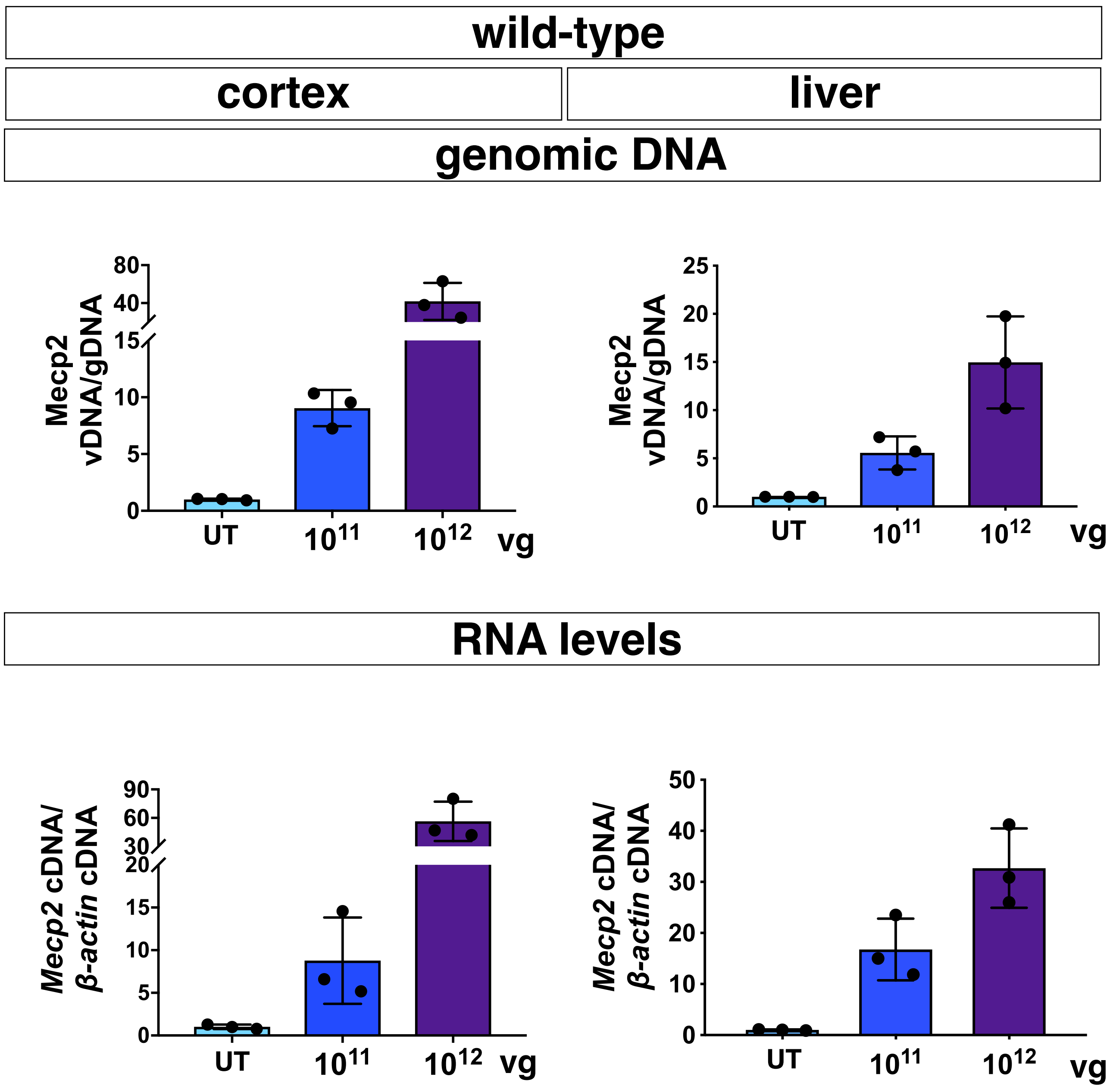

### Supplementary Figure 8

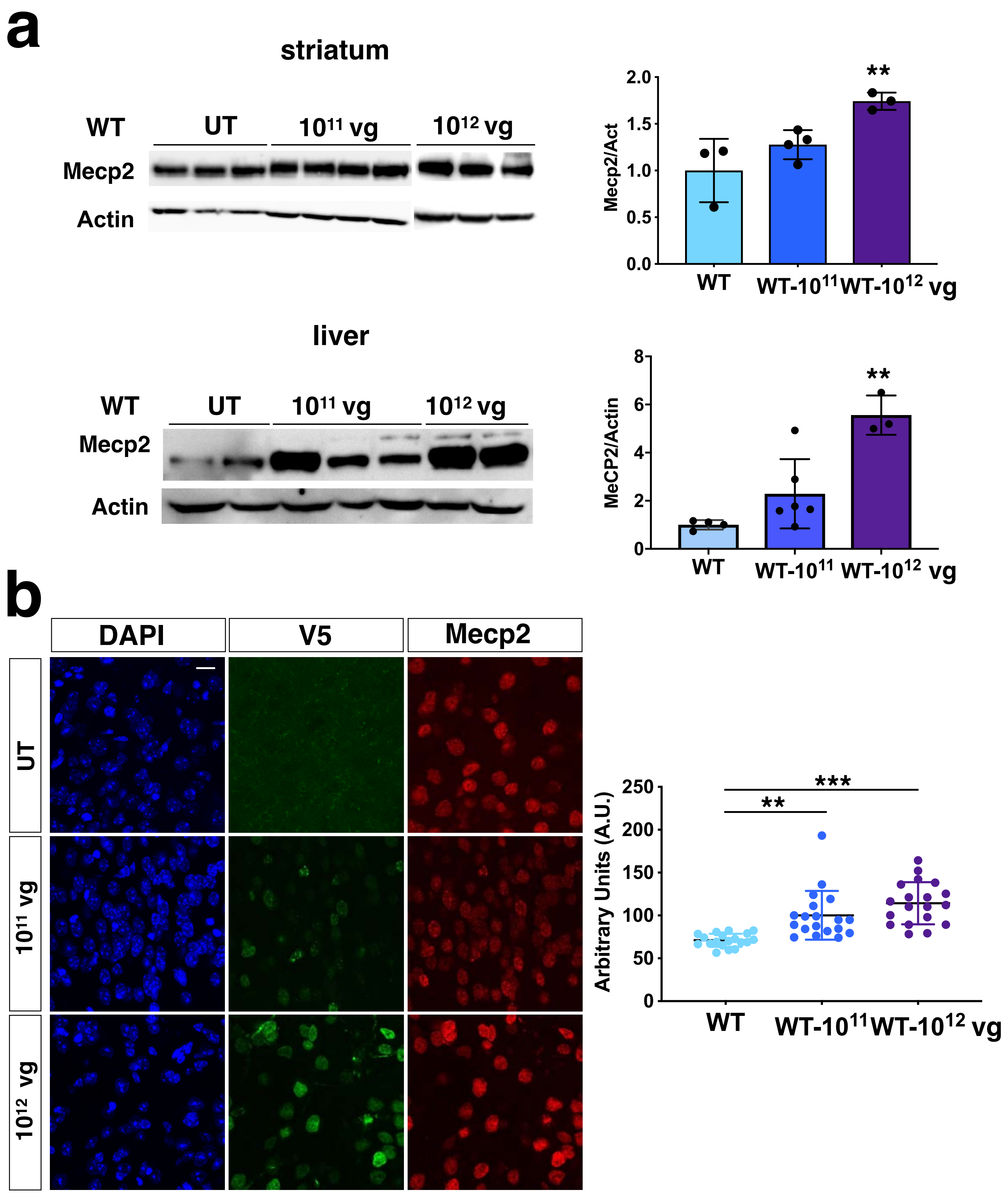

**Supplementary movie**

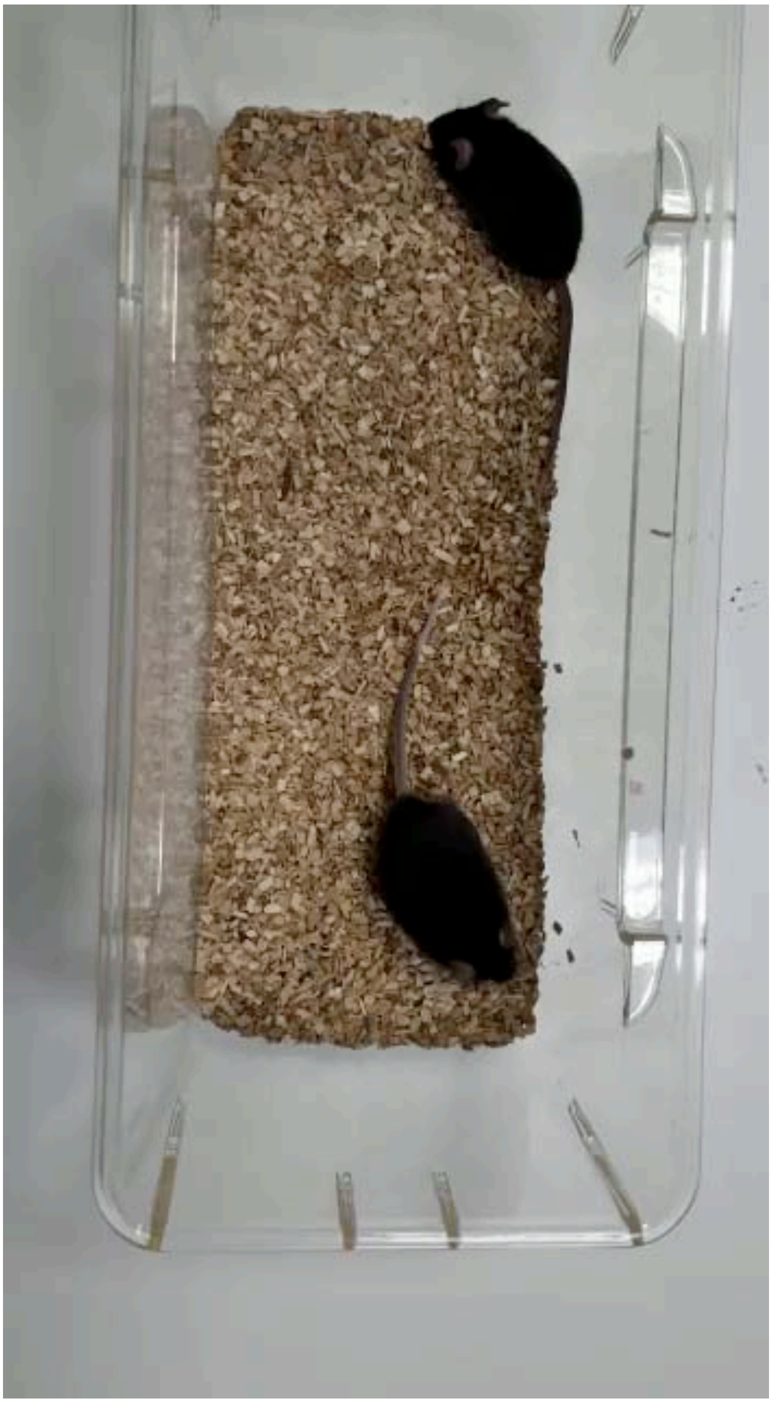

| **Mice** | **Wild-Type** | | **Mecp2$^{-/y}$** | | | **Reference value #** | **Normal Range** |
|---|---|---|---|---|---|---|---|
| **AAV (dose)** | None | iMecp2 ($10^{12}$) | None | GFP ($10^{12}$) | iMecp2 ($10^{12}$) | None | |
| **ALB (g/dL)** | 3,23 ± 0,39 | 3,50 ± 0,69 | 3,17 ± 0,75 | 3,43 ± 0,64 | 3,85 ± 0,35 | 3,87 ± 0,20 | 2,7 / 3,6 |
| **ALP (U/L)** | 109 ± 54 | 116 ± 18 | 131 ± 14 | 114 ± 28 | 140 ± 22 | 140 ± 16 | 100 / 140 |
| **ALT (U/L)** | 43 ± 11 | 52 ± 27 | 71 ± 2 | 60 ± 6 | 68 ± 13 | 79 ± 19 | 0 / 70 |
| **Liver HE** | 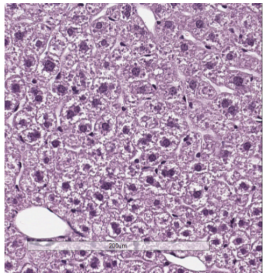 | 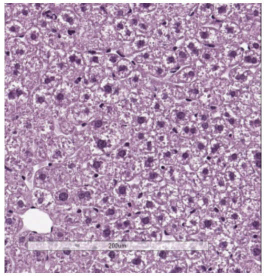 | 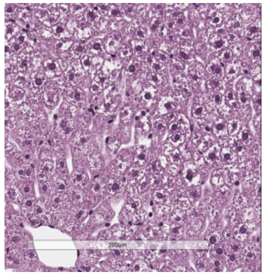 | 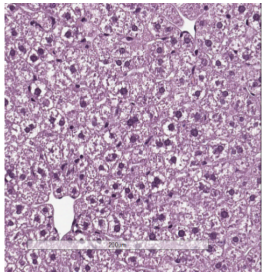 | 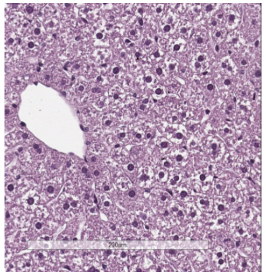 | | |

**Supplementary Table 1**: No evidence of liver toxicity in mice administered with high dose of AAV treatments. Blood serum levels of liver enzyme and liver histochemical analysis (representative images) were used as indicators of liver health. # Reference values for C57BL/6J male mice were taken from the mouse phenome database https://phenome.jax.org/. Abbreviations, Treat: treatment, ALB: albumin, ALP: Alkaline phosphatase, ALT: Alanine aminotransferase, HE: hematoxylin/eosin staining. All values are indicated as mean ± SD, n = 3, SD. Scale bar: 200 µm

å

**Supplementary Table 2**

Primers employed for qRT-PCRs.

| Name | Sequence |
|---|---|
| *Mecp2*-cDNA-endo-F | 5′- ggaggtgtgatgcaggagaa-3′ |
| *Mecp2*-cDNA-endo-R | 5′- tttggattgggttgatgtgg-3′ |
| *Mecp2*-cDNA-ecto-F | 5′-gggagtagttggagcattgg-3′ |
| *Mecp2*-cDNA-ecto-R | 5′-agggatgccacccgtagat-3′ |
| *Mecp2*-cDNA-tot-F | 5′-ccggggacctatgtatgatg-3′ |
| *Mecp2*-cDNA-tot-R | 5′-aagcttttccctggggatt-3′ |
| *β-Actin*-F | 5′-tggcaccacaccttctacaat-3′ |
| *β-Actin*-R | 5′-aggcatacagggacagcaca-3′ |
| 18S-F | 5′- gtaacccgttgaaccccatt-3′ |
| 18S-R | 5′-ccatccaatcggtagtagcg-3′ |
| *Sqle*-F | 5′-aaccaaccaagtgcagagtg-3′ |
| *Sqle*-R | 5′-ccgattacagcatcatcttcc-3′ |
| *Nsdhl*-F | 5′-acatggtggagcagttgctg-3′ |
| *Nsdhl*-R | 5′-cctttgagagctgggtacaggt-3′ |
| *MsmoI*-F | 5′-tgggctgtgcagtcattgagg-3′ |
| *MsmoI*-R | 5′-atggagcctgaaactcgtgat-3′ |
| *Kcnj10*-F | 5′-cgtcggtcgctaaggtctat-3′ |
| *Kcnj10*-R | 5′-gcaatgtgctccattctcac-3′ |
| *Kcnc3*-F | 5′-ggaccgagcttgcttccttg-3′ |
| *Kcnc3*-R | 5′-cgttggcgttgaggtcgg-3′ |
| *Angpl4*-F | 5′-caacgccacccacttaca-3′ |
| *Angpl4*-R | 5′-aatcactgtccagcctccat-3′ |
| *Mecp2*-gDNA-F | 5′-agaaagcctgggagtgtggt -3′ |
| *Mecp2*-gDNA-R | 5′-cttccttgacctcgatgctg -3′ |
| *Lmnb2*-gDNA-F | 5′-gttaacactcaggcgcatgggcc-3′ |
| *Lmnb2*-gDNA-R | 5′-ccatcagggtcacctctggttcc-3′ |

**Supplementary Table 3**

**List of antibodies for flow cytometry.**

| **Reactivity** | **Fluorophore** | **Clone** | **Company** |
|---|---|---|---|
| Mouse | PE-CD3 | 17A2 | BD Pharmingen |
| Mouse | PE-Cy7-CD4 | GK1.5 | E-Bioscence |
| Mouse | FITC-CD8a | 53-6.7 | BD Pharmingen |
| Mouse | APC-Cy7-CD44 | IM7 | BD Pharmingen |
| Mouse | BV786-CD62l | Mel-14 | BD Pharmingen |
